# Genetically-Encoded Fluorescence Barcodes Allow for Single-Cell Analysis via Spectral Flow Cytometry

**DOI:** 10.1101/2024.10.23.619855

**Authors:** Xiaoming Lu, Daniel J. Pritko, Megan E. Abravanel, Jonah R. Huggins, Oluwaferanmi Ogunleye, Tirthankar Biswas, Katia C. Ashy, Semaj K. Woods, Mariclaire W.T. Livingston, Mark A. Blenner, Marc R. Birtwistle

## Abstract

Genetically-encoded, single-cell barcodes are broadly useful for experimental tasks such as lineage tracing or genetic screens. For such applications, a barcode library would ideally have high diversity (many unique barcodes), non-destructive identification (repeated measurements in the same cells or population), and fast, inexpensive readout (many cells and conditions). Current nucleic acid barcoding methods generate high diversity but require destructive and slow/expensive readout, and current fluorescence barcoding methods are non-destructive, fast, and inexpensive to readout but lack high diversity. We recently proposed theory for how fluorescent protein combinations may generate a high-diversity barcode library with non-destructive, fast and inexpensive identification. Here, we present an initial experimental proof-of-concept by generating a library of ∼150 barcodes from two-way combinations of 18 fluorescent proteins, 61 of which are tested experimentally. We use a pooled cloning strategy to generate a barcode library that is validated to contain every possible combination of the 18 fluorescent proteins. Experimental results using single mammalian cells and spectral flow cytometry demonstrate excellent classification performance of individual fluorescent proteins, with the exception of mTFP1, and of most evaluated barcodes, with many true positive rates >99%. The library is compatible with genetic screening for hundreds of genes (or gene pairs) and lineage tracing hundreds of clones. This work lays a foundation for greater diversity libraries (potentially ∼10^5^ and more) generated from hundreds of spectrally-resolvable tandem fluorescent protein probes.

## Introduction

Cell barcoding, a collection of methods by which single cells (or sometimes populations) are uniquely labeled, has been and remains fundamental to answering a wide variety of biological questions^1–15^. Applications include studying cell type and/or state dynamics (e.g. evolution of tumor heterogeneity^15–18^), tracking lineage (e.g. cell fate decisions^4,14,15,19–21^), or identifying genetic perturbations in a cell (e.g. CRISPR screening)^12,22–27^. Barcodes are usually made from either fluorophores (small molecules or fluorescent proteins) or nucleic acids, with each having benefits and drawbacks depending on the questions at hand^1–4,6,7,23,26–35^. In general, an ideal barcode library has high diversity (many unique barcodes), is fast/inexpensive to readout (analyze many cells and conditions), and its observation is non-destructive (repeated measurements of the same cell or population).

Fluorescent barcodes can be generated from combinations of fluorophores (fluorescent proteins or small molecules), with additional diversity sometimes possible from an intensity dimension^1,3,4,20,28,36–38^. Fluorescent barcodes are usually non-destructive/fast/inexpensive to readout via flow cytometry or microscopy. However, they to-date lack high diversity relative to nucleic acid approaches or genome scale. Small molecule fluorophore approaches have not yet been used to identify genetic perturbations^1,3,28^. Fluorescent protein (FP) barcodes are genetically-encoded but have even less diversity^4,20,37–39^. Alternatively, nucleic acid barcodes are usually random DNA sequences^26,27,29–35,40^, but are sometimes more structured^41,42^, and have incredible diversity (reviewed in^15,21,43^). For example, just 10 bases can generate ∼10^6^ (4^10^) barcodes. However, the readout is often single-cell (or bulk) sequencing, which is destructive and often much slower and more expensive than flow cytometry or microscopy. These properties prevent observation of the cell after barcode identification and hinder large-scale experiments, although have the benefit of often providing the transcriptome^1,22^. Thus, an open question therefore is whether FP-based approaches may be scaled, given some of their advantages relative to nucleic-acid-based approaches, and if so how.

We recently proposed theory for and simulation studies supporting a fluorescent protein-based, single-cell barcoding method that bridges fast, non-destructive fluorescence readouts with the large barcode diversity^44^. The barcoding approach is based on Multiplexing using Spectral Imaging and Combinatorics (MuSIC)^28,44–46^, which generates unique emission spectra signatures from combinations of individual fluorescent proteins. In this paper, we demonstrate initial experimental proof-of-principle by generating ∼150 barcodes from 18 fluorescent proteins, with experimental testing of 61. We use a pooled cloning strategy to generate a barcode library that is validated to contain every possible combination of the 18 fluorescent proteins. Experimental results using single mammalian cells and spectral flow cytometry demonstrate excellent classification performance of individual fluorescent proteins, with the exception of mTFP1, and of most evaluated barcodes, with many true positive rates >99%. The library is compatible with genetic screening for hundreds of genes (or gene pairs) and lineage tracing hundreds of clones. This work lays a foundation for greater diversity libraries (potentially 10^5^ and more) generated from hundreds of spectrally-resolvable tandem fluorescent protein probes.

## Results

### MuSIC Barcodes in Single Cells

While previous work explored the concept of a MuSIC barcode and their application to high-dimensional single cell analysis^44^, experimental demonstration remained. To answer the question of whether MuSIC barcodes could be constructed, characterized, and then reliably identified in single cells, we targeted a simple application consisting of two fluorescent protein (FP) barcodes. While this application is a small fraction of the theoretical potential, we reasoned it nevertheless provides a sizable library diversity for some biological applications and would support larger scale efforts if successful.

Specifically, we selected 18 FPs spanning UV-to-IR spectral properties (EBFP2^47^, mTagBFP2^48^, mT-Sapphire^49^, mAmetrine^49^, mCerulean3^50^, LSSmOrange^51^, mBeRFP^52^, mTFP1^53^, EGFP^54^, CyOFP1^55^, mClover3^56^, mVenus^57^, mPapaya^58^, mOrange2^59^, mRuby3^56^, mKate2^60^, mCardinal^61^, miRFP670^62^). These are called “probes”, in this case simply individual FPs (**Fig. 1A**). A MuSIC barcode, in this study, is a combination of two probes. Given 18 FPs and barcodes made of 2 probes, 153 unique barcodes can be generated. It is important to note here we only count combinations, where order does not matter, as opposed to permutations, where order matters. That is because in the targeted application using single-cell fluorescence emission spectra measurements, the order of the barcode would not be distinguishable. Furthermore, we consider combinations without replacement, that is, we do not consider a barcode consisting of two of the same FP. While these may exist in the eventual construction, we suspect inclusion of such barcodes will deteriorate detection reliability. That is because if only a single FP is identified, it may be due to a false negative for the 2^nd^ FP.

**Figure 1.**
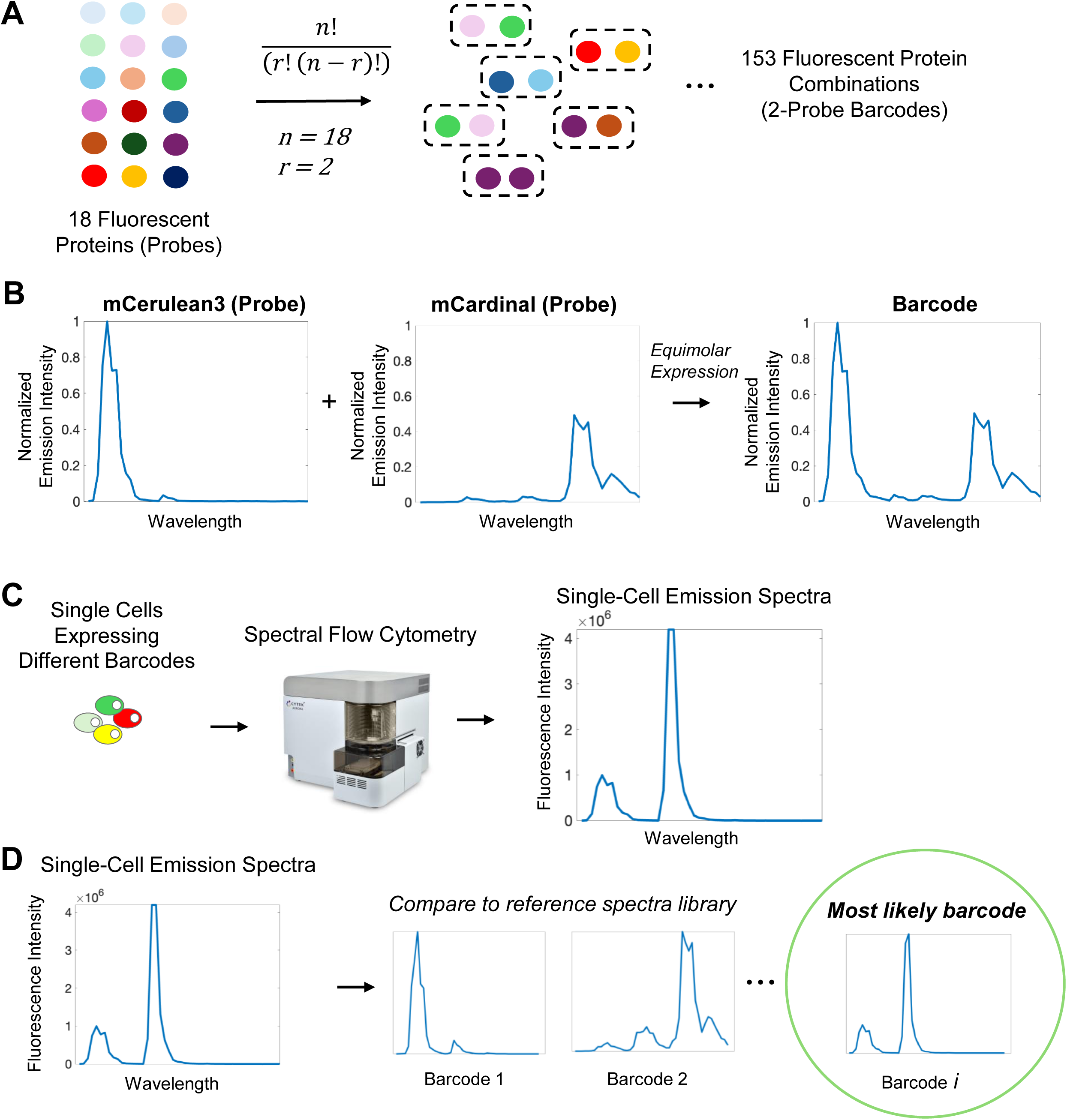
Barcodes and Their Analysis. **(A)** 153, 2-probe barcodes can be generated from 18 fluorescent proteins (FPs). Combinations (vs permutations) are relevant because FP order does not affect the barcode. **(B)** Barcodes have a unique emission spectra arising from the two FPs. HEK293T cells were transfected and individual FP spectra measured by spectral flow cytometry. The barcode spectra is illustrative as the sum of the two FP spectra. **(C)** Spectral flow cytometry measures emission spectra of individual cells with potentially different barcodes. **(D)** The emission spectra for each cell can then be compared to a reference library (containing spectra for individual FPs and each barcode) to identify which barcode the cell mostly likely possesses.

As an example, consider a cell expressing an mCerulean3 and mCardinal barcode (**Fig. 1B**). The individual FP spectra are invariant to whether a barcode is mCerulean3 / mCardinal or mCardinal / mCerulean3. The combination of these individual spectra gives the barcode spectra, balanced by their relative expression levels (in this example equimolar), which in principle are unique from all the other barcodes. This uniqueness, of course, depends on the spectral emission detector properties. If different single cells in a population are expressing different barcodes, full spectrum flow cytometry can be used to read out single-cell fluorescence emission spectra (**Fig. 1C**). These measured emission spectra can be compared to known references, from which the most likely barcode can be identified (**Fig. 1D**). This conceptual setup forms the basis of the experimental testing of the approach in what follows.

### Construction of a MuSIC Barcode Library

As mentioned above, MuSIC barcodes are constructed here by combining two fluorescent proteins (although they could be larger). To accomplish this, we generated two separate sets of probes (**Fig. 2A**). Each set has its own backbone, either pReceiver (pR) or pMulti (pM). Every fluorescent protein was cloned into each backbone. To subsequently generate barcodes, we took a pooled construction approach, via Golden Gate assembly with BbsI (**Fig. 2B**). For efficiency of design, barcode expression is driven by a single promoter in pReceiver, and the barcode elements are separated by 2A peptide sequences. We found that only a single T2A sequence led to some fusion protein generation (which would generate “confusing” spectra), and that a tandem P2A-T2A (tPT2A) sequence^63^ was necessary to make such fusion events largely negligible (**Fig. S1**). We did not test a single P2A sequence, so it is possible that a single P2A could also make fusion events negligible. Each of the 18 plasmids in both sets were pooled and then digested. The digested pR and pM pools were combined in equimolar amounts and a ligation was performed, forming a pooled barcode library. We note that this approach is designed to scale to larger barcodes but for proof-of-concept purposes here we analyze barcodes with two fluorescent proteins.

**Figure 2.**
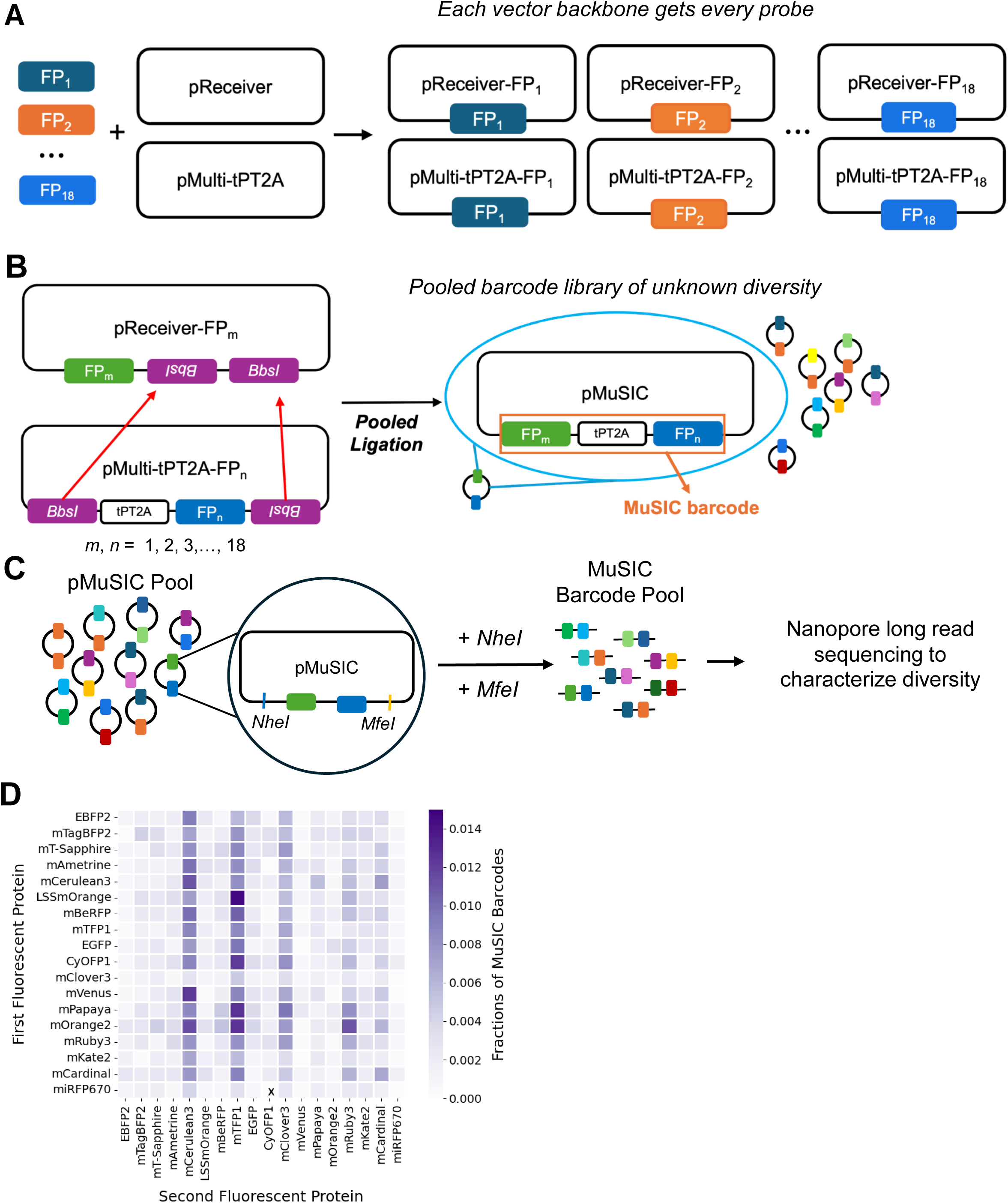
Barcode Library Construction and Validation. **(A)** Each of the 18 fluorescent proteins (FPs) were cloned into the pReceiver and pMulti backbones. **(B)** The FPs from pMulti were inserted into the pReceiver backbone by GoldenGate assembly in a pooled format to generate a pMuSIC barcode library. **(C)** The pMuSICs were double-digested with *NheI* and *MfeI* and then sequenced using a nanopore long read sequencer (MK-1C). This avoids PCR and short-read alignment which may be problematic due to FP homology. **(D)** Nanopore sequencing data from the pMuSIC pool were analyzed and individual FPs in each barcode assigned and counted. Heatmap color denotes the percentage of reads attributed to a particular FP in either the 1st or 2nd position of the barcode. We verified that all potential barcodes are represented in the pool, except for miRFP670-CyOFP1, as indicated by the marked ‘x’. However, this barcode was found during single colony isolation (**Table S16**).

The proportion of each barcode type in the pooled library was unknown, so we performed sequencing (**Fig. 2C**). When considering the type of sequencing to use, we had to address two main concerns. One was that we had to know the correct pairing of the fluorescent proteins of each barcode. Short read sequencing may not provide enough data to determine such pairing, so we decided to use long read sequencing. The next concern was that some fluorescent proteins have high sequence homology, which can complicate PCR^64,65^. Nanopore sequencing became apparent as the best option for our purpose as it would allow us to perform long read sequencing while also eliminating the need for PCR that is required by most other deep sequencing technologies. To analyze the sequencing results, we determined which fluorescent protein was in the first position, and which was in the second (see Methods and Code). While there appears to be bias towards mCerulean3, mTFP1, and mClover3 in the second position, observable due to the deeper color in those columns (possibly due to systematic pipetting error occurring during the pooling/digestion stage), all barcode combinations (with the exception of one subsequently found during single colony isolation below) were found to be present, as indicated by color spread throughout the heatmap visualization (**Fig. 2D**). These results validate the cloning process used to generate the MuSIC barcode pool and provide a plasmid library ready for testing the approach in cells.

### Unmixing of Single Fluorescent Proteins and Generating Reference Spectra

Before testing the identification of barcodes, we wanted to ensure that each of the 18 FPs were deconvolvable from one another, while also generating the reference spectra for each FP needed for the identification of barcodes. To test whether the 18 FPs could be unmixed, we individually transfected each of the 18 pR-FP plasmids and assayed cells with a spectral flow cytometer. This experiment was performed at least twice independently to test whether the emission spectra were stable and reproducible, with results showing near uniform concordance (**Fig. S2**). We repeated this experiment in a different cell line (MDA-MB-231 versus HEK293T), and the spectra were highly concordant (**Fig. S3**).

The analysis strategy (**Fig. 3A-B**) partitioned all positively transfected cells into training and testing pools (done in triplicate). To classify which of the 18 FPs was most likely present in a single cell, the data from (48) individual fluorescent channels were unmixed, the unmixed levels (19; 18 FPs plus 1 autofluorescence; analogous to a relative concentration) were compared to a threshold (estimated from training) to compute a score, and the maximum score was chosen to finish classification. During training, the threshold for each FP was estimated from the pooled data one-at-a-time, by sliding a classification threshold and computing a ROC curve, with the chosen threshold being that which maximized true positives while minimizing false positives (**Fig. S4**).

**Figure 3.**
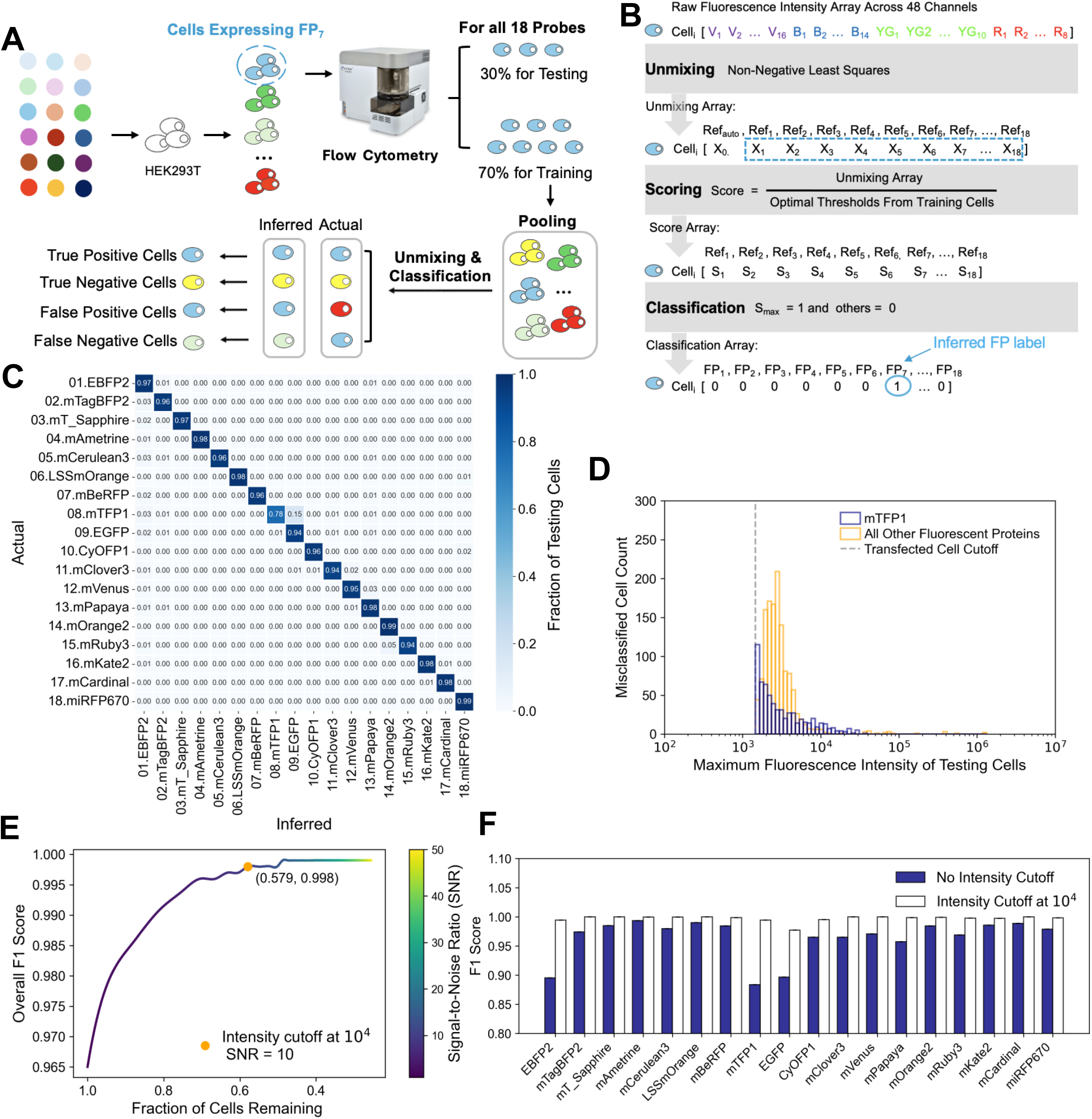
Single-Cell Identification of Individual Fluorescent Proteins. **(A)** Plasmids encoding individual fluorescent proteins (FPs) were transfected and analyzed by spectral flow cytometry. Three random splits of 70% for training and the remaining for testing were used. All cells from all 18 FPs were pooled for training, testing and analysis. **(B)** Data from a single cell are unmixed, scored for each FP by comparing to a training-estimated threshold, and then classified based on the maximum score. **(C)** Average inferred percentage of the testing pool cells (n=3) for each FP, for each transfected sample. Actual is the transfected FP; inferred is the classified FP. **(D)** The number of misclassified cells as a function of their maximum fluorescence intensity. Cells with low intensity are predominantly the misclassified ones, and mTFP1-containing cells are the most frequently misclassified. **(E)** Average overall F1 scores (n=3) were calculated after eliminating cells with a maximum fluorescence intensity below the cutoff. We proceeded with the cutoff as indicated (10^4^; SNR=10). **(F)** F1 score for each FP with and without the intensity cutoff (10^4^).

To evaluate performance on the testing set, we first analyzed the fraction of cells with correct (diagonal) and incorrect (off-diagonal) labels, stratified by FP (**Fig. 3C**). Overall, a large proportion of cells had correct labels (dark blue on the diagonal). The most notable deviation is mTFP1 (8^th^ row and column), which was often misclassified as EGFP and accounted for a substantial fraction of all classification errors. More subtly, most FPs were prone to be misclassified as EBFP2 (1^st^ column).

We reasoned that low fluorescence intensity of a single cell may also be a factor in misclassification. The distribution of misclassification frequency as a function of signal intensity demonstrates that cells with intensity lower than 10^4^ (relative intensity) are responsible for most errors, and that again mTFP1 is responsible for a disproportionate fraction of errors (**Fig. 3D**).

To further evaluate performance, we focused on the F1 score, which balances multiple dimensions of the classification problem (true positives, false positives, and false negatives), and the effect of an intensity cutoff suggested by the above (**Fig. 3E-F**). Although an intensity cutoff may improve performance, there is a competing objective of reducing the size of the dataset by eliminating cells. We found that an intensity cutoff of 10^4^, which is a signal-to-noise ratio (SNR) of ∼10 (given background at ∼10^3^), substantially improves the F1 score of the overall dataset (which considers all FPs simultaneously), while retaining more than half of the cells (**Fig. 3E**). Thus, moving forward in this work we retain this cutoff, but stress that different applications may consider a different weighting. F1 score stratified by individual FPs (**Fig. 3F**) shows the chosen intensity cutoff substantially improves performance for mTFP1, EGFP, and EBFP2, which were the most problematic FPs noted above. We conclude that individual FPs can be reliably classified, and a single-cell fluorescence intensity cutoff will be applied (∼10^4^) to improve classification performance.

### Identification of MuSIC Barcodes in Single Cells

To assess the validity of MuSIC barcode identification in single cells, we decided to isolate individual barcodes so precisely controlled experiments could be performed (**Fig. 4A**). To do this, we performed repeated single-colony pickup following transformation of the pooled library, and then determined which barcodes were found by sequencing (**Fig. S5**). We obtained 112 pMuSIC plasmids, of which 107 were identified as valid MuSIC barcodes, while 5 with identical fluorescent proteins (e.g. EGFP-EGFP) were classified as invalid. Out of these, 70 were unique permutations, and 61 were unique combinations. We transfected cells with single MuSIC barcodes and analyzed them with spectral flow cytometry (**Fig. 4B**).

**Figure 4.**
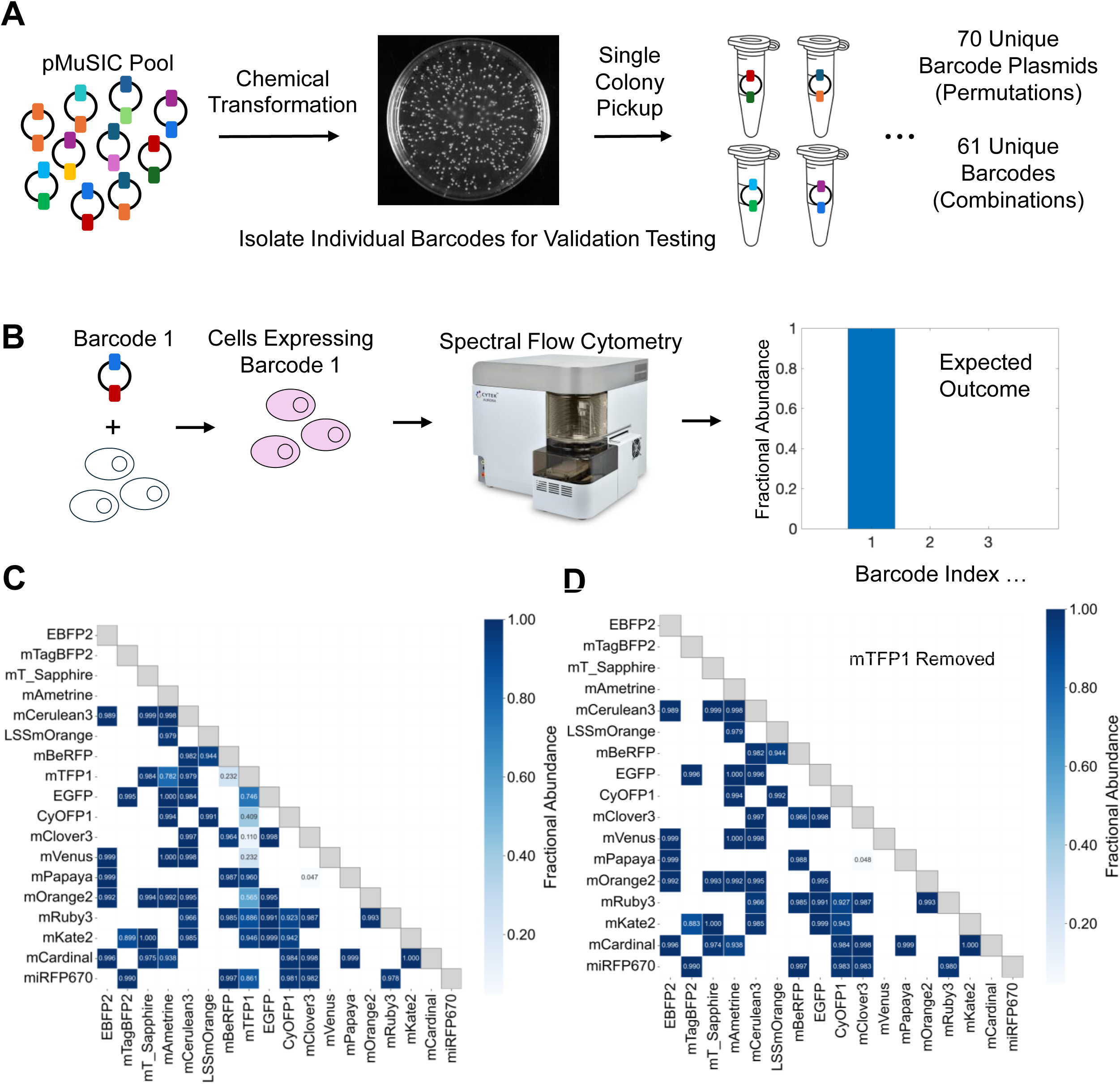
Single-Cell Barcode Identification. **(A)** Single-colony pickup after pool transformation enables isolation of individual barcode plasmids (further detail in Fig. S5) **(B)** Single barcodes are transfected and then analyzed to assess barcode identification performance by fractional abundance. **(C-D)** Fractional abundance for barcodes isolated and assayed as in A-B, with (C) or without (D) inclusion of mTFP1-containing barcodes. White squares denote barcodes that were not isolated and analyzed.

Overall, barcodes could be identified with high fidelity (**Fig. 4C**). However, it was clear that mTFP1-containing barcodes did not perform well, despite the applied intensity cutoff. Therefore, we removed mTFP1-containing barcodes, which demonstrated improved performance (**Fig. 4D**). One exception was the combination of mClover3 and mPapaya, which was incorrectly identified in nearly every cell. Further, mTagBFP2/mKate2 and CyOFP1/mRuby3 had reduced performance. The CyOFP1/mRuby3 barcode was particularly interesting because we isolated both permutations (CyOFP1-mRuby3 and mRuby3-CyOFP1), and one had very good performance (0.998—mRuby3-CyOFP1) whereas the other did not (CyOFP1-mRuby3—0.859).

The spectra of the individual fluorescent proteins and of the expected versus measured barcodes (**Fig. S6**) reveal a possible shared explanation for the reduced performance of these barcodes. In theory, each of these barcodes would be uniquely resolvable, based on expected spectra (**Fig. S6**). However, the expression of the first fluorescent protein in the barcode is much lower than expected, which follows from the difference between the expected spectra and the measured spectra essentially missing the first fluorescent protein characteristics (**Fig. S6A-C**). This gives rise to a measured spectra from which it is difficult to infer the correct barcode, since the spectra looks essentially only like the second fluorescent protein. The fact that the alternative, good performing permutation mRuby3-CyOFP1 shows larger than expected contribution of CyOFP1 to the measured barcode spectra (**Fig. 6B**) is consistent with this interpretation. Also, if this were true, then expected spectra for barcodes with good performance (**Fig. S6D-F**) would be closer to the measured spectra. Two examples show close correspondence between the two spectra (mKate2-mCardinal and EGFP-mAmetrine) whereas one does not (mAmetrine-mVenus). For the good performing barcode with substantial deviation between expected and measured spectra (mAmetrine-mVenus), the reduced intensity of the first FP remains well above the noise level, which we expect gives rise to the retained ability for accurate identification. Why only some barcodes display this issue remains unclear. One possibility is cell-line specificity. However, the spectra of these barcodes are nearly identical in MDA-MB-231 cells (**Fig. S7**), implying that is unlikely the reason. It may be due to artifacts of tPT2A cleavage, particularly because in the cases explored, the first FP is affected. It may also be related to burden, which is now understood to potentially be significant in mammalian systems^66,67^. Future studies will be needed to understand the underlying mechanism. We conclude that overall, MuSIC barcodes can be identified with excellent performance in cells using spectral flow cytometry, with a few exceptions.

## Discussion

Cell barcoding is a core technique in the biological sciences. Barcodes are typically made from either nucleic acids or fluorophores. Ideally, the barcoding method would enable (i) a large number of unique barcodes to be constructed (i.e. library size or diversity) such that many single clones or populations could be analyzed simultaneously, (ii) non-destructive readout to enable tracking the same cells or population over time, and (iii) fast and/or inexpensive readout to enable the same but additionally measurement across many conditions. Historically, nucleic acid methods excel at (i) but struggle with (ii) and (iii), whereas fluorescence methods, the opposite. In this work, we establish a proof-of-concept that combinations of fluorophores and spectral emission measurements increase potential library size for fluorescence-based measurements. We create a library of 153 barcodes from 2-way combinations of 18 fluorescent proteins, and demonstrate that many can be reliably identified with true positive rates of >99% using spectral flow cytometry.

Because this library is genetically-encoded, it could potentially be coupled to small gRNA libraries for fast single-cell genetic or genetic interaction screening, or for lineage tracing of hundreds of clones. In terms of genetic interaction screening, current methods based on incorporation of two guide RNAs in a single plasmid^25,27^ are fundamentally limited by the number of individual bacteria that can contain a unique plasmid in the library. Our approach may not be as limited, because a single gRNA could be coupled to a single barcode. Spectral live-cell microscopy imaging to trace clones in space and time could enable analysis of an even greater number of clones, since the likelihood of observing the same barcode in the same spot is lower, and thus the same barcode may be repeated in different spatial locations given appropriate experimental design. Compared to nucleic-acid methods, however, we do not envision our approach contributing to doublet or multiplet discrimination as recent work has shown^68^, especially since widely-used light-scattering discrimination exists for flow cytometry.

The established library size here, while somewhat greater than the potential of current fluorescent protein approaches^4,36,37^, remains far below that possible using nucleic acids^26,27,29–35^. We envision multiple routes to higher library diversity. One could be simply more advanced demultiplexing and classification algorithms, which could allow for more accurate detection of fluorescent proteins and larger barcodes. Here, we employed a somewhat simple approach based on classical linear unmixing (non-negative least squares), followed by selection of the two fluorescent proteins most likely to be present. The continuous estimated expression level from linear unmixing was converted to a ratio score by comparison to a threshold established from analysis of single fluorescent proteins. The two fluorescent proteins (because it is a two-member barcode) with the highest scores were then said to comprise a cell’s inferred barcode, thus binarizing the output for casting as a classification problem. We expect modern machine learning methods could further improve the already good classification performance we observed^69,70^. Alternatively, exploiting expression level as a barcode dimension could greatly increase diversity. However, given the observed strong effect of fluorescence intensity cutoffs on classification performance, natural cell-to-cell variability in protein expression^71^, as well as the finite range of intensity detection in the spectral flow cytometer, using expression level as a barcode dimension may prove challenging.

While differences in classification methods could impact effective library size and performance, the largest potential increases come from modifying the number of fluorescent proteins, and the number of fluorescent proteins per barcode. This proof-of-concept work is based our recent theory and simulations^44^, that considers not only individual fluorescent proteins, but fluorescent protein fusions as independent potential barcode members. The approach, called multiplexing using spectral imaging and combinatorics (MuSIC)^45^, works because when a fluorescent protein fusion is constructed, if substantial FRET occurs from one to the other, then the emission spectra of the fusion protein cannot be constructed from a linear combination of the two individual fluorescent proteins, and is thus unique in terms of linear unmixing. Based on the same 18 fluorescent proteins used here, we expect that hundreds of so-called MuSIC probes could be constructed using 2 and 3-way fusion proteins, which has some technical precedent^72–74^. Simulations matched to current spectral flow cytometry equipment suggested that using only 2-probe barcodes, from a pool of hundreds of barcodes, could enable human genome scale library diversity (∼10^4^ – 10^5^)^44^. If one could make 3-probe barcodes from 200 probes, the potential diversity becomes ∼10^6^. Future work will explore these avenues.

Another potential improvement relates to hardware. Emission spectra scanning has historically been possible but technically far more challenging than simple filter-based four color measurements. However, spectral flow cytometers have recently enabled much greater multiplexing than previously possible^75,76^. White lasers that enable excitation wavelength tuning are now more common^77^, as are microscopes with spectral detection^78,79^. All such hardware advances are expected to enhance feasibility of reading out these emission spectral barcodes and to increase the fidelity of classification for larger libraries. In this work, we found that mTFP1 was problematic in our panel, but it was initially unclear whether this was due to hardware limitations, or due to features unique to the HEK293T cell line used. The fact that spectra for individual FPs (**Fig. S3**) were nearly identical in another cell line (MDA-MB-231), implies mTFP1 difficulties may be overcome by different hardware. Both good and difficult barcodes similarly had nearly identical spectra in another cell line (**Fig. S7**), which implies some poor barcode performance could also be improved with different hardware. <u>However, some, as noted above, may be an inherent property of the barcode itself which requires more work to understand fully.</u> Thus, improving performance and scaling to more barcodes will likely involve hardware/instrumentation changes, but further testing of biological scenarios remains important.

There are notable limitations of the approach, some of which derive from sequence homology among many fluorescent proteins. First is how to stably deliver such barcodes to cells. Many barcoding approaches use pooled lentiviral libraries, but creating lentiviral libraries with fluorescent protein combinations is likely to be problematic due to template switching from sequence homology^31,33^. Silent mutations (and perhaps other non-critical non-synonymous mutations) could be used to potentially overcome such issues^80^, although such variants would need careful testing because silent mutations can affect expression through mRNA folding^81^. Delivery to cells may be enabled by simple transfection coupled with recombination-based landing pad systems for control of single copy integration in safe harbors through selection and counter-selection^82^. Barcode library construction may be hampered by difficulty performing PCR on segments and constructs containing large homologous regions from fluorescent proteins. Indeed, here we avoided PCR of intact barcodes because of such concerns, which led us to nanopore sequencing-based characterization of the pooled barcode library. Although recombinase-deficient *E. coli* strains are routinely used, recombination of barcodes may be a problem during library propagation or in the eventual target cells-of-interest. However, the success of Brainbow systems *in vivo* argues this may achievable^4,36,37^. As the size of probes and barcodes grow, the limits of plasmid length may be reached. Recent work has highlighted expression burden may be a significant phenomenon even in mammalian systems^66,67^, and the extent to which it impacts the current manifestation or future implementations is as yet unclear. Lastly, we removed mTFP1 from analysis here as our particular 4-laser spectral flow cytometer struggled to accurately identify barcodes containing it. The fact that individual FP spectra were nearly identical in another cell line implies the mTFP1 difficulties are mostly likely a result of hardware limitations. However, we suspect a 5-laser model may be successful for such purposes, and clearly the particulars of which fluorescent proteins can be demultiplexed are highly dependent upon the excitation channels used and the emission spectra detection properties.

In conclusion, we demonstrate here construction of a library of ∼150 fluorescence barcodes from combinations of 18 fluorescent proteins, and experimental classification of a subset using spectral flow cytometry. There is substantial room for growth in library size via both design and hardware. We expect that this approach could fulfill a gap in barcoding technology to provide a large diversity library with fast, inexpensive and non-destructive readout. This could enable large-scale, single-cell genetic and genetic interaction screening or lineage tracing applications.

## Methods

### Experimental

#### Availability

All final plasmids were verified by sequencing (*Genewiz* or *Plasmidsaurus*). They are available on Addgene (pool—#227193; individual plasmids—#228594-228631).

#### Molecular Cloning

##### Preparation of Chemically Competent Cells

One microliter of DH5α from NEB 5-alpha Competent *E. coli* (*New England Biolabs*, Cat# C2987I) was inoculated in 2.5 mL SOB medium (*Fisher BioReagents*, Cat# BP9737-500) in a 14 mL round-bottom test tube (*Corning*, *Falcon Cat#* 352059), incubated at 225 rpm and 37 °C overnight. The overnight culture was then diluted 1:1,000 (v/v) into 200 mL SOB medium and incubated at 37 °C at 225 rpm until the OD_600_ reached 0.32 (about 5 hours). After chilling on ice for 10 minutes, cells were centrifuged at 1000 x *g* (∼3000 rpm) for 10 minutes at 4 °C with a Fiberlite F15-8x50c rotor (*Thermo Scientific*) in a Sorvall Legend XFR refrigerated centrifuge (*Thermo Scientific*) and gently resuspended in 16 mL ice-cold CCDB80 transformation buffer. This buffer was prepared using 10 mM potassium acetate (*VWR Chemicals BDH*, Cat# 9254-500G), 80 mM calcium chloride dihydrate (*Sigma-Aldrich*, Cat# C3306-100G), 20 mM manganese chloride tetrahydrate (*Alfa Aesar*, Cat# 44442-25G), 10 mM magnesium chloride hexahydrate (S*igma-Aldrich*, Cat# 13152-1KG), 10% glycerol v/v (*ICN Biomedicals*, Cat# 800687), and pH adjusted to 6.4 by 0.1 N hydrochloric acid (*VWR Chemicals BDH,* Cat# BDH3028-2.5LG). After chilling the cells on ice for another 20 minutes, they were centrifuged again at 1000 x *g* for 10 minutes at 4 °C. The cell pellet was gently resuspended in sterile ice-cold CCMB80 buffer to achieve a final OD_600_ of ∼1. Cells were aliquoted (50 μL) into pre-chilled sterile microcentrifuge tubes and frozen in a dry ice/ethanol bath for 5 minutes before long-term storage at -80 °C. Transformation efficiency with pUC19 Control DNA at different dilution ratios *(New England Biolabs,* Cat# N3041A) showed ∼7.4 x 10^7^ cfu/µg DNA.

##### Chemical Transformation

A 50 µL aliquot of lab-prepared DH5α competent cells, or, in some cases, NEB 5-alpha Competent *E. coli* (*New England Biolabs*, Cat# C2987I) were thawed on ice for 5-10 minutes. One microliter (unless otherwise noted) of ligation (or other) product was added to the cells, and the tube was carefully flicked 5 times to mix the DNA and competent cells. After 30 minutes incubation on ice, the cells were heat-shocked at 42 °C for exactly 30 seconds, followed by immediate transfer back to ice for 5 minutes. Then, 950 µL of room temperature SOC outgrowth medium (*New England Biolabs, Cat#* B9020S) was added to the cells in a 5 mL polypropylene round-bottom tube (*Corning*, *Falcon Cat#* 352063). The mixture was shaken at 225 rpm at 37 °C for 1 hour and around 100 µL was then spread onto a selection LB agar plate. The plates (*Fisher Scientific*, Cat# FB0875712) were made from LB Agar Lennox (*Fisher BioReagents*, Cat# BP9745-500) according to the manufacturer’s instruction and supplemented with 50 µg/mL Kanamycin (*VWR*, Cat# 0408-10G) for pReceiver backbone or 25 µg/mL Chloramphenicol (*ACROS Organics*, Cat# AC227920250) for pMulti backbone.

##### Inoculation

Around 2.5 µL glycerol stock or a single colony was inoculated into 2.5 mL autoclaved LB medium supplemented with appropriate antibiotics in a 14 mL round-bottom test tube (*Corning*, *Falcon* Cat# 352059), incubating at 225 rpm, 37°C overnight. LB medium was prepared by first combining 5 g Tryptone (*Fisher Bioreagents*, Cat# BP1421-500), 2.5 g Yeast Extract, Bacteriological Grade (*VWR Life Science*, Cat# J850-500), and 5 g Sodium Chloride (*Fisher BioReagents*, Cat# BP358-1) in 400 mL of Milli-Q water (Millipore Advantage A10 water purification system). The pH was adjusted to 7.0 with 10 N Sodium Hydroxide (*Fisher Chemical*, Cat# SS255-1) and then Milli-Q water added to a final volume of 500 mL.

##### Colony Polymerase Chain Reaction (PCR)

Single colonies were screened by colony PCR with screening primers (*Integrated DNA Technologies, IDT*) as shown in **Table S1**. For each test, approximately 0.2 µL of overnight culture was added to the reaction mixture containing 1.5 µL 10 x *Taq* ThermoPol® buffer (*New England Biolabs*, Cat# B9004S), 0.3 µL 10 mM dNTP mix (*Thermo Scientific*, Cat# R0192), 0.2 µL of 20 µM forward primer, 0.2 µL of 20 µM reverse primer, 0.1 µL *Taq* DNA polymerase (*New England Biolabs*, Cat# M0267L), and autoclaved Milli-Q water to a final volume of 15 µL. The PCR was initiated at 95 °C for 5 minutes to release the DNA. A series of 10 touchdown cycles were then done, consisting of 95 °C for 30 sec, starting 65 °C and decreasing 1 °C per cycle for 30 sec, followed by 68 °C for 30 sec. An additional 25 PCR cycles with an annealing temperature at 52 °C was performed, followed by a final extension at 68 °C for 5 minutes. Candidates were selected by performing electrophoresis on a 1% (g/100 mL) agarose gel (*Fisher BioReagents*, Cat# BP160-500) prepared in 1 x TAE buffer, which was diluted from 50 x TAE buffer. To prepare 50 x TAE buffer we combine 121 g Tris (*Thermo Scientific*, Cat# J22675-A1), 28.55 mL Glacial Acetic Acid (*VWR BDH Chemicals*, Cat# BDH3098-3.8LP), and 50 mL 0.5 M Ethylenediaminetetraacetic acid pH 8.0 (EDTA; *Alfa Aesar*, Cat# A10713), and add Milli-Q water to a final volume of 500 mL.

##### Restriction Digests

For cloning purposes, approximately 1 µg of DNA (plasmid or PCR product) was added to a digestion mixture containing 5 µL of rCutSmart Buffer (New England Biolabs, Cat# B6004S), 10-20 units of each restriction enzyme, and autoclaved Milli-Q water, all having a final volume of 50 µL. The reaction mixture was incubated at 37 °C overnight to achieve maximum digestion efficiency.

For verification purposes, colony PCR products were confirmed by single or double digestion with restriction enzymes as shown in **Table S2**. For each reaction, approximately 0.5 µg DNA was added into the digestion mixture containing 1.5 µL rCutSmart™ Buffer (*New England Biolabs*, Cat# B6004S), 10 units of each restriction enzyme, and autoclaved Milli-Q water, all having a final volume of 15 µL. The reaction mixture was incubated at 37 °C for at least 4 hours.

##### Backbone Construction (Fig. S8)

Different resistance genes were used for pReceiver (pR) and pMulti (pM). All sequences of synthetic DNA fragments used for construction of these backbones are listed in **Table S3**. All PCR was performed using the primers listed in **Table S4** and Q5 High-Fidelity DNA Polymerases (*New England Biolabs*, Cat# M0491L), following the manufacturer’s instructions. Plasmids were purified by the PureYield miniprep system (*Promega,* Cat# A1222).

For the pR backbone, the plasmid was divided into three parts: the f1 origin (f1ori), Kanamycin resistance cassette, and the CMV promoter. The f1ori was synthesized by *IDT* and then further amplified. The other two parts were similarly amplified from mTFP1-N1 (*Addgene*, #54521). Amplicons from synthesized DNA fragments were purified using the Monarch PCR & DNA Cleanup Kit (New England Biolabs, Cat# T1030L), while amplicons from plasmids were purified using the Monarch Gel DNA Extraction Kit (New England Biolabs, Cat# T1020L). The pR plasmid was then created with these purified PCR products using Gibson Assembly Master Mix (*New England Biolabs*, Cat# E2611L) with a 1:1:1 molar ratio and incubated at 50 °C for 1 h.

For the pR-NheI backbone, an NheI site was inserted before the CMV enhancer by overlap extension PCR of the SnaBI-ApaLI double-digested fragment to create the SnaBI-*NheI*-ApaLI insert. This was inserted into the pR backbone through SnaBI (*New England Biolabs,* Cat# R0130S) and ApaLI (*New England Biolabs,* Cat# R0507S) digest followed by ligation with *T4* ligase (*New England Biolabs,* Cat# M0202L).

For the pM backbone, the plasmid was also divided into three parts: the T2A-f1ori, Chloramphenicol (CMR) resistance cassette, and the CMV promoter. The T2A-f1ori and CMR cassettes were synthesized by *IDT* and *Genewiz*, respectively. The CMV cassette was amplified and purified as above. The pM plasmid, designed to contain the *BbsI-T2A-BsaI-spacer-BsaI-BbsI* cassette, was constructed using these purified PCR products through Gibson assembly as described above.

For the pM-tPT2A backbone, a tandem P2A-T2A(tPT2A) DNA fragment was synthesized by *IDT* and then amplified and purified as described above. Both purified tPT2A amplicons and the pM backbone plasmids were treated with BbsI-HF (*New England Biolabs*, Cat# R3539L) to create the insert and the vector, respectively. They were purified with a Select-a-Size DNA Clean & Concentrator kit (*Zymo Research*, Cat# D4080) and then ligated by Goldengate assembly with an insert-to-vector ratio as 3:1 and the *T4* ligase (*New England Biolabs,* Cat# M0202L), following the manufacturer’s instructions.

##### Cloning Individual Fluorescent Proteins into Different Backbones

The workflow for inserting individual fluorescent proteins into pR, pR-NheI, pM, and pM-tPT2A is illustrated in **Fig. S9A** (gray). Sequences of fluorescent proteins, listed in **Table S5**, were designed to avoid recognition sites of type IIS restriction enzymes, including BsaI, Esp3I, and BbsI, through the implementation of silent mutations. They generated from direct synthesis and amplified with Q5 High-Fidelity DNA Polymerases (*New England Biolabs*, Cat# M0491L), using primers according to **Fig. S9B** and **S9C** and **Table S6**. For reverse primers longer than 60nt (**Fig. S9B**), each reverse primer was split into two shorter ones. After two rounds of PCR, the final amplicons containing the BsaI-FP-TAA-BsaI cassette were prepared as inserts. They were then purified and subsequently inserted into backbones through BsaI sites using Goldengate assembly with *T4* ligase (*New England Biolabs,* Cat# M0202L) and BsaI-HF (*New England Biolabs*, Cat# R3733L).

##### Cloning mTFP1-mVenus Fusion

To compare the cleavage efficiency of T2A versus tPT2A in MuSIC barcodes, we cloned the mTFP1-mVenus fusion into pM backbone to create a positive-control probe for poorly cleaved MuSIC barcodes. The mTFP1 and mVenus were linked with seven amino acids (AGGGGLG), as described previously^45^. The pM backbone was digested with BsaI-HF (*New England Biolabs*, Cat# R3733L) overnight. At the same time, mTFP1 was amplified with primers as listed in **Table S7** to generate the insert (mTFP1-Esp3I-TAA-Esp3I), allowing the mVenus to be inserted through the Esp3I sites. Both the first insert and the vector were purified and ligated by Goldengate assembly with the BsaI-HF (*New England Biolabs*, Cat# R3733L) and *T4* ligase (*New England Biolabs,* Cat# M0202L), following the manufacturer’s instructions. The resulting plasmid (pM-mTFP1-Esp3I-TAA-Esp3I) was screened and verified as above. This plasmid was then used for the insertion of a second fragment (linker-mVenus), which was amplified from the pQLinkHD-mVenus plasmid (*Addgene*, #118861)^45^. Both the second insert and vector were purified and ligated by Goldengate assembly with Esp3I (*New England Biolabs*, Cat# R0734L) and *T4* ligase (*New England Biolabs,* Cat# M0202L), following the manufacturer’s instructions.

##### Construction of MuSIC Barcodes for 2A Cleavage Efficiency Comparison

Both pR-FPs and pM-FPs were digested overnight with BbsI-HF (*New England Biolabs*, Cat# R3539L). The digestion products from the pR probes were purified with the Monarch PCR & DNA Cleanup Kit (New England Biolabs, Cat# T1030L) to serve as the vector, while those from pM probes (T2A-FP) were isolated using the DNA Gel Extraction Kit (New England Biolabs, Cat# T1020L) to serve as the insert. The purified vector and insert were then mixed at a molar ratio of 1:5 (vector to insert) and combined by Goldengate assembly through BbsI cutting sites to generate pR-FP-T2A-FP as the pMuSIC-T2A constructs for cleavage efficiency comparison (**Fig. S9A**). Similarly, pR-NheI-FP and pM-tPT2A-FP were digested and purified prior to Goldengate assembly. These were then used to create pR-NheI-fp-tPT2A-FP as the pMuSIC-tPT2A constructs for cleavage efficiency comparison (**Fig. S9A**).

##### Construction of MuSIC Barcode Pool

(**Figs. S5 and S9**) First, 0.45 pmol of each pR-NheI-FP and pM-tPT2A-FP were pooled to create two separate pR and pM probe libraries, respectively. For pM-mCardinal, T2A was used (as opposed to tPT2A), but this did not affect downstream classification performance to our knowledge. Each of these libraries were then digested separately overnight using BbsI. The pR probe pool was purified using a Monarch PCR & DNA cleanup kit, while the digested pM probe pool was purified using the DNA Gel Extraction Kit (New England Biolabs, Cat# T1020L). To prevent self-ligation, the pR probe pool was treated with Quick CIP (*New England Biolabs,* Cat# M0525S) and purified using a DNA cleanup kit as described above. The pR and pM probe pools were assembled at a 1:5 vector-to-insert molar ratio using *T4* ligase (*New England Biolabs,* Cat# M0202L) and BbsI-HF (*New England Biolabs*, Cat# R3539L). A ligation reaction without the insert (pM probe pool) was also set up as the negative control group, which was verified to have negligible colonies post-transformation. The assembly products were diluted at 1:10 and 1:100 ratios with autoclaved Milli-Q water, and 1 μL used to transform DH5α chemically competent cells. For each dilution in each group, 100 µL of the transformation culture were spread onto an LB agar/Kana+ plate A while the rest were onto a second plate B. After incubation for 21 hours at 35 °C, the colonies were imaged using a ChemiDoc™ XRS+ imager (*Bio-Rad*).

##### Isolation of Single MuSIC Barcodes

Single colonies were selected for barcode isolation (**Fig. S5**). Plasmid DNA was extracted using a PureYield miniprep system (*Promega,* Cat# A1222) and further confirmed by double digestion. Barcode identity was determined by sequencing (*Plasmidsaurus*). Barcodes containing identical FPs were discarded.

##### Pooled MuSIC Barcode Library Generation

Transformed colonies were scraped from the agar plate into 10 mL of LB medium supplemented with 50 µg/mL kanamycin (*VWR*, Cat# 0408-10G) in a sterile 50 mL polypropylene conical tube (*Corning*, *Falcon* Cat# 352070) using a cell scraper (*Fisher Scientific*, Cat# 08100-241). The cell scraper was washed with 4 mL of LB/Kana+ twice, while the agar plate was gently washed with 2 mL of LB/Kana+. The resulting cell/medium mixture was carefully transferred into the conical tube by pipetting to maximize the transfer of colonies into the tube. The cell/medium mixture was then added to 170 mL of LB/Kana+ in a 1 L flask and incubated at 250 rpm at 37 °C for 18 hours until O.D.600 reached 3, as measured by a NanoDrop™ 2000 spectrophotometer (*Thermo Scientific*). The pooled pMuSIC library was isolated by the PureYield™ plasmid maxiprep system (*Promega*, Cat# A2393).

#### Cell Culture

HEK293T cells were obtained from ATCC (CRL-3216) and the MDA-MB-231 cell line (ATCC, Cat# HTB-26) was kindly provided by Dr. Adam Melvin from Clemson University. Both cell lines were grown in Dulbecco’s Modified Eagle’s Medium (DMEM) (*Gibco*, Cat# 10313021) supplemented with 10% (v/v) fetal bovine serum (*Gibco*, Cat# 10082139) and 2mM L-Glutamine (*Corning*, Cat# 25-005-CI) under a 5% CO_2_ atmosphere at 37 °C. HEK293T cells were sub-cultured every 2-3 days, while MDA-MB-231 cells were sub-cultured every 3-4 days to prevent confluence. This process involved lifting with Trypsin (*Gibco*, Cat# 25200-072), followed by centrifugation at 100 x *g* (∼1000 rpm) for 5 minutes at room temperature, and a final resuspension in full growth medium (as described above).

#### Transfection

For flow cytometry experiments, HEK293T cells (6 x 10^4^) and MDA-MB-231 cells (7-10 x 10^4^) were seeded onto 12-well plates (*Corning*, *Falcon* Cat# 353043) 18-24 hours before transfection. Once cells reached approximately 20-30% confluency, they were transfected with pR or pM or pMuSIC plasmids at concentrations ranging from 50 ng to 300 ng for HEK293T cells and 300 ng to 500 ng for MDA-MB-231 cells using *Lipofectamine 3000* (*Invitrogen,* Cat# L3000-008) according to the manufacturer’s instructions (**Table S10**). For higher transfection efficiency, the medium for both cell lines was replaced with DMEM/L-glutamine one hour prior to transfection. Additionally, HEK293T cells were supplemented with 10% FBS, while the medium for MDA-MB-231 cells was replaced with fresh growth medium four to six hours after transfection.

For western blotting, HEK293T cells (5 x 10^5^) were seeded into 6-well plates (*Corning*, *Falcon* Cat# 353046) and incubated for 24 h to achieve ∼50% confluency. Three micrograms of DNA were transfected into each well by Lipofectamine 3000 (*Thermo Fisher Scientific*, Cat# L3000001).

#### Western Blotting

Forty-eight hours post-transfection, cells were lysed in 100 µL of fresh, ice-cold RIPA buffer per well^83^. The plates were kept on ice for 15 minutes and the cells were agitated every 5 minutes to ensure thorough lysis. The lysates were scraped off with cell scrapers (*Stellar Scientific*, Cat# TC-CS-25) and transferred into pre-chilled 1.5 mL microcentrifuge tubes. Each tube was vortexed for 5 sec 3 times to homogenize cell debris. Cells were then centrifuged at 4 °C at 16,000 x *g* for 5 minutes to clear the lysate.

About 10 µL supernatant was stored at -80 °C or directly used for protein quantification with a Pierce Rapid Gold BCA Protein Assay kit (*Thermo Scientific*, Cat# A53225) according to the manufacturer’s instructions. The standard curve was generated using Bovine Serum Albumin Standard Ampules (*Thermo Scientific*, Cat# 23209), and the concentration of the samples was determined with a BioTek *SynergyH1* multimode microplate reader. The rest of the supernatant was transferred into a new pre-chilled 1.5 mL micro-centrifuge tube and mixed with 4 x Laemmli sample buffer (*Bio-Rad*, Cat# 161-0747) that had been supplemented with 10% 2-Mercaptoethanol (*Fisher Scientific*, Cat# O3446I-100) for sample preparation. The tubes were then heated at 95 °C for 5 minutes in a dry block heater and stored at -80 °C.

Thirty micrograms of protein were loaded into wells along with a Chameleon Duo Pre-Stained Protein Ladder (*LICOR*, Cat# 928-60000). The proteins were separated by a Tris (*Thermo Scientific*, Cat# J22675-A1) - glycine (*Thermo Scientific*, Cat# A13816.0E) SDS gel, with all ingredients listed in **Table S8** in the Mini PROTEAN Tetra Cell (*Bio-Rad*, Cat# 165-8001). The protein samples were subsequently transferred onto a 0.2 µm Immun-Blot PVDF Membrane (*Bio-Rad*, Cat# 162-0177) in a glycine/methanol (*Fisher Chemical*, Cat# A452SK-4) transfer buffer at 100V for 1 hour. The membrane was blocked with 0.1% (v/v) Tween-20 (*Fisher Scientific*, Cat# BP337-100) in Tris-buffered saline (TBST) supplemented with a final concentration of 5% BSA w/v (*Fisher Scientific*, Cat# BP1600-100) for 1 h at room temperature and then incubated with the primary antibody solution with gentle rocking on a BioRocker 2D rocker (*Denville Scientific INC*) at 4 °C overnight. After washing three-times with TBST buffer at room temperature, 15 minutes each, the membrane was incubated with the secondary antibody solution while gently rocking on a rotator mixer stirrer (*Fisher Scientific 2309FS*) at room temperature for 30 minutes. The membrane was washed again as previously described (but without Tween) and the target protein bands were visualized by a LI-COR Odyssey infrared fluorescence scanning system.

Immunoblotting was performed using antibodies diluted as specified in **Table S9**. All primary antibodies were diluted as above in TBST supplemented with a final concentration of 5% BSA w/v and 0.02% Sodium Azide w/v (*Thermo Scientific Chemicals,* Cat# 190380050) while the secondary antibodies were diluted in 1x TBST buffer supplemented with a final concentration of 0.1% SDS (*Fisher Bioreagents*, Cat# BP2436-1) and 0.02% Sodium Azide w/v.

#### Flow Cytometry

Forty-eight hours post-transfection, cells were lifted using 0.25% Trypsin (*Gibco*, Cat# 25200-072) and centrifuged at 300 x *g* for 5 minutes at 4 °C. Cell pellets were then washed with 1 mL of pre-chilled phosphate-buffered saline (PBS) (*Fisher BioReagents*, Cat# BP2944-100) and centrifuged as above. The cells were resuspended in 0.5 mL ice-cold FACS buffer (PBS, 1% BSA w/v) for analysis using a spectral flow cytometer (Cytek Aurora). Cells transfected with pR backbone were used as a negative control.

Cells were gated based on FSC and SSC, and singlets were gated by FSC-H versus FSC-A. Cytek Aurora settings are initially configured for the use of SpectroFlo QC Beads (*Cytek*, Cat# B7-10001), which are ∼3 µm in diameter, Lot 2004 for HEK293T cells and Lot 2005 for MDA-MB-231 cells. These settings are not optimal for analyzing HEK293T cells (11 to 15 µm in diameter), nor for MDA-MB-231 cells (8 to 12 µm in diameter). To account for this size difference, the area scaling factors (ASFs) and gains needed to be adjusted to make sure the fluorescence intensity was accurate and on scale. To maintain the accuracy of the fluorescence intensity, the height (H) and area (A) intensities needed to be equivalent at the peak channel of each cell. For example, the peak channel of mRuby3 is YG1, therefore the YG area scaling factor was adjusted until the fluorescent intensities of YG1-A and YG1-H were roughly the same. We repeated this for each of the eighteen pR-FP probes, adjusting the area scaling factors relative to the peak channel as needed. Since different fluorescent proteins with varying intensity levels were transfected at different DNA amounts on both cell lines(**Table S10**), the gains were adjusted to ensure that the fluorescence remained on scale. This adjustment was made until the brightest sample did not cause overflow in the peak channel. All the final instrument setting parameters for pR-FP references are listed in **Table S11**. The normalized median fluorescence intensity (MFI) of the manually gated positive population for each pR-FP sample across 48 channels (**Table S12**) was used to generate the spectra of the corresponding FP probe (**Fig. S2**). We verified that so long as the area scaling factors and gains remained constant, the FP spectra were constant in replicates.

#### Nanopore Sequencing

##### Analysis of pR-FP Pool

Approximately 4.5 picomoles of each pR-FP plasmid (equivalent to 1.05 µg of DNA per 50 µL reaction, across 10 reactions) were first treated with NheI-HF (*New England Biolabs*, Cat# R3131L) overnight, followed by an additional overnight digestion with MfeI-HF (*New England Biolabs*, Cat# R3589L) to achieve maximum digestion efficiency. After heat inactivation at 80 °C for 20 minutes, all ten reactions of double-digested pR-FP fragments were pooled into a 2 mL-DNA LoBind Eppendorf tube and supplemented with 1/10 v/v of sodium acetate (3 M, pH 5.2) (*Thermo Scientific*, Cat# R1181), 0.05 µg/µL of Glycogen (Thermo Scientific, Cat# R0561), and 2.5 volumes of absolute ethanol (*Fisher Scientific*, Cat# BP2818-4), and concentrated via ethanol preciptation. All tubes were inverted gently until fully mixed and incubated at –20 °C overnight. After centrifugation at 16,000 x *g* for 30 minutes at 4 °C, the DNA pellets were washed with ice-cold 70% ethanol and centrifuged again for 15 minutes at 4 °C. They were then air-dried for 10 minutes at room temperature and dissolved in nuclease-free water. The concentrated fragments (NheI-CMV-FP-MfeI) were subsequently isolated using the Monarch Gel DNA Extraction Kit (New England Biolabs, Cat# T1020L). The concentration of each purified probe was measured by a Qubit Flex fluorometer (*Invitrogen by Thermo Fisher Scientific*) using a Qubit 1X dsDNA HS assay kit (*Invitrogen*, Cat# Q33231).

Approximately 250 fmol of each digested fragment was subjected to end-prep with a NEBNext Ultra II End repair/dA-tailing Module (*New England Biolabs*, Cat# E7546S). They were then purified with the AMPure XP Beads provided from the native barcoding kit 24 (V14) (*Oxford Nanopore*, Cat# SQK-NBD114.24) and quantified via Qubit. Each probe was ligated with native barcodes from the nanopore native barcoding kit using the blunt/TA ligase master mix (*New England Biolabs*, Cat# M0367S). The ligation products from individual FPs were pooled and underwent a second round of purification using the AMPure XP Beads, followed by another Qubit quantification. The pooled probes were further ligated with the adapters from the native barcoding kit using the NEB quick T4 ligase (*New England Biolabs*, Cat# E6056S). After a third round of purification and quantification, approximately 20 fmol of barcoded FP library was prepared and loaded into a MinION flow cell (*Oxford Nanopore*, Cat# R10.4.1), which was then inserted into a MinION MK1C device (*Oxford Nanopore*) for nanopore sequencing.

##### Analysis of Known MuSIC Barcodes

Twenty pMuSICs verified by sequencing (*Plasmidsaurus*) were selected for native-barcoded nanopore sequencing as shown in **Table S13**. Approximately 1.9 picomoles of each pMuSIC (equivalent to 1 µg of DNA per 50 µL reaction, across 3 reactions) were double digested with NheI-HF (*New England Biolabs*, Cat# R3131L) and MfeI-HF (*New England Biolabs*, Cat# R3589L) overnight. Each pMuSIC was end-prepped and ligated with a native barcode provided from the native barcoding kit 24 (V14) (*Oxford Nanopore*, Cat# SQK-NBD114.24) and mixed prior to the adaptor ligation for the subsequent native-barcoded nanopore sequencing, as described above. Approximately 20 fmol of barcoded library was loaded into a MinION flow cell (*Oxford Nanopore*, Cat# R10.4.1), which was then inserted into a MinION MK1C device (*Oxford Nanopore*) for nanopore sequencing.

##### Analysis of MuSIC Barcode Pool

The MuSIC barcodes pool prepared by maxiprep was double-digested using 1 µg of DNA per 50 µL reaction, across 10 reactions, with NheI-HF (*New England Biolabs*, Cat# R3131L) and MfeI-HF (*New England Biolabs*, Cat# R3589L) overnight. The ligated products were heat inactivated at 80 °C for 10 minutes and concentrated by ethanol precipitation for electrophoresis. DNA fragments (NheI-CMV-fp-tPT2A-fp-MfeI) containing MuSIC barcodes were isolated by the Monarch Gel DNA Extraction Kit (New England Biolabs, Cat# T1020L) and subject to end-prep, adapter ligation, and DNA cleanup and quantification as described above to prepare the library for nanopore sequencing using a ligation sequencing kit V14 (*Oxford Nanopore*, Cat# SQK-LSK114). Approximately 20 fmol of the pooled MuSIC barcode library was prepared and loaded into a MinION flow cell (*Oxford Nanopore*, Cat# R10.4.1), which was then inserted into a MinION MK1C device (*Oxford Nanopore*) for nanopore sequencing.

### Computational

#### Availability

The code and data used in this study are available upon request, and the code is publicly available on github (github.com/birtwistlelab/MuSIC-Fluorescent-Protein- Barcodes--Nanopore_Sequencing_Analysis and MuSIC-Fluorescent-Protein-Barcodes-- Spectral_Unmixing).

#### Nanopore Sequencing

First reads are filtered, which consists of two steps. Only reads that passed internal Nanopore quality checks and were in the expected length range (1.2 – 1.8 kb for pR-FP; 2.0 to 2.8 kb for known barcodes; and 2.0 to 3.0 kb for unknown barcodes) were retained. Additionally, sequences were aligned and reoriented to the expected 5’->3’ direction with the CMV promoter at the beginning using *Align.PairwiseAligner* in global alignment mode, and only sequences with over 80% alignment to the CMV sequence were retained. Next, reads are scored against all reference sequences (single FPs or barcodes). For each read, the highest alignment value for each FP or barcode (18 versus 324 possibilities—considering order is necessary for sequence-level analysis) was assigned a 1, and all other 0. This classification could then be compared to known labels for the 18 single FP or 20 barcode controls, which revealed excellent performance (**Fig. S10**), and gives confidence for assignment in cases of unknown barcodes (**Fig. 2**). Tables S15-17 contain additional information regarding these results.

#### Fluorescent Protein and Barcode Classification (Fig. 3A-B)

##### Data Export

Flow cytometry (FCS) files, containing fluorescence intensities of each cell across 48 channels from the Cytek Aurora, were converted to CSV files with scale values using FlowJo v10.10.0 (*BD Life Science*). Only positively transfected cells were analyzed, and the gating threshold for defining positive versus negative cells was based on the population transfected with empty vector, usually with a cutoff ∼10^3^ [AU] in the maximum fluorescence intensity channel for a particular FP.

##### Reference Spectra for Single FPs

A subset of the positively transfected cell population with moderate fluorescence intensity for each FP was selected to determine the reference spectra, calculated from the median fluorescence intensity for each of the 48 channels minus that of the negatively transfected cells for the same. The autofluorescence spectra was also estimated, giving a total of 19 reference spectra.

##### Classification

Classification of which FP (or two FPs) is in a single cell consists of three steps. First, a positively transfected cell is unmixed by non-negative least squares, using the reference spectra from above and the intensity profile for that cell. This transforms a 48 x 1 vector to a 19 x 1 vector of estimated relative abundances. Next, each relative abundance is divided by a respective threshold estimated from training (see below), giving an 18 x 1 vector of scores for each FP in that cell. Finally, the FP (or two FPs) with the highest score (or scores) are classified to be present in that cell, and the others absent.

##### Parsing Data into Testing and Training

All single FP-transfected cells were randomly assigned to training or testing sets, in a 70/30 split. Three different random partitions were generated and results averaged across them.

##### Estimating Thresholds from Training Data

For each FP-of-interest, training cells from all 18 FP transfections were used for estimation of that FPs threshold. At a given threshold, for a particular FP, all cells with unmixed relative abundance greater than the threshold are classified as positive, and those below as negative. For the FP-of-interest, comparison to the known labels enables calculation of the number of true positives and false negatives. For the other FPs, comparison to the known labels enables calculation of the number of true negatives and false positives. By repeating this procedure for several thresholds, a ROC curve could be constructed for each FP-of-interest (**Fig. S4**). The chosen threshold was that closest to the (0,1) point on the ROC curve.

## Supporting Information

There are supplementary figures and tables associated with this manuscript.

## Acknowledgements

This work was supported by NIH grant R35GM141891 to MRB. DJP received funding from the Department of Education Grant P200A180076. We thank the Clemson Creative Inquiry program for undergraduate research funding, and the Clemson University Departments of Chemistry, Chemical and Biomolecular Engineering, and Biomedical Data Sciences and Informatics for graduate student support.

## Author contributions

Xiaoming Lu: conceptualization; methodology; investigation; formal analysis; data curation; writing-original draft; writing-review and editing.

Daniel J. Pritko: conceptualization; investigation; formal analysis; writing-original draft.

Megan E. Abravanel: investigation; data curation.

Jonah R. Huggins: data curation.

Tirthankar Biswas: investigation.

Katia C. Ashy: investigation.

Oluwaferanmi Ogunleye: investigation.

Mariclaire W.T. Livingston: investigation.

Semaj K. Woods: investigation.

Mark A. Blenner: conceptualization; writing-review and editing.

Marc R. Birtwistle: conceptualization; methodology; data curation; funding acquisition; supervision; visualization; writing-original draft; writing-review and editing.

## Table of Contents Graphic

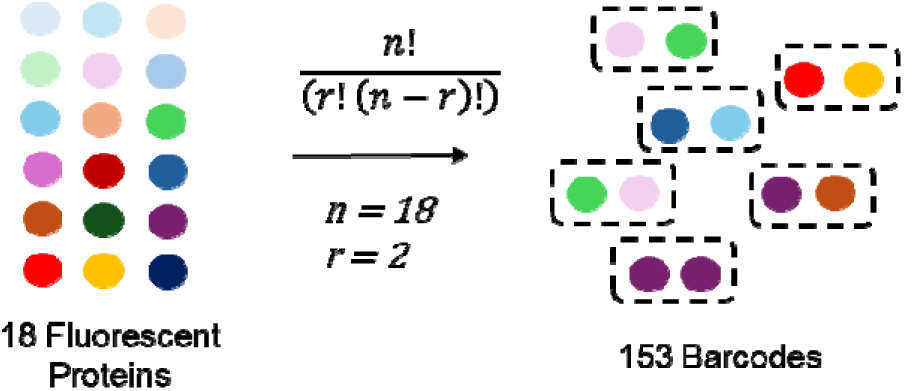

## Supporting Information

**Figure S1.**
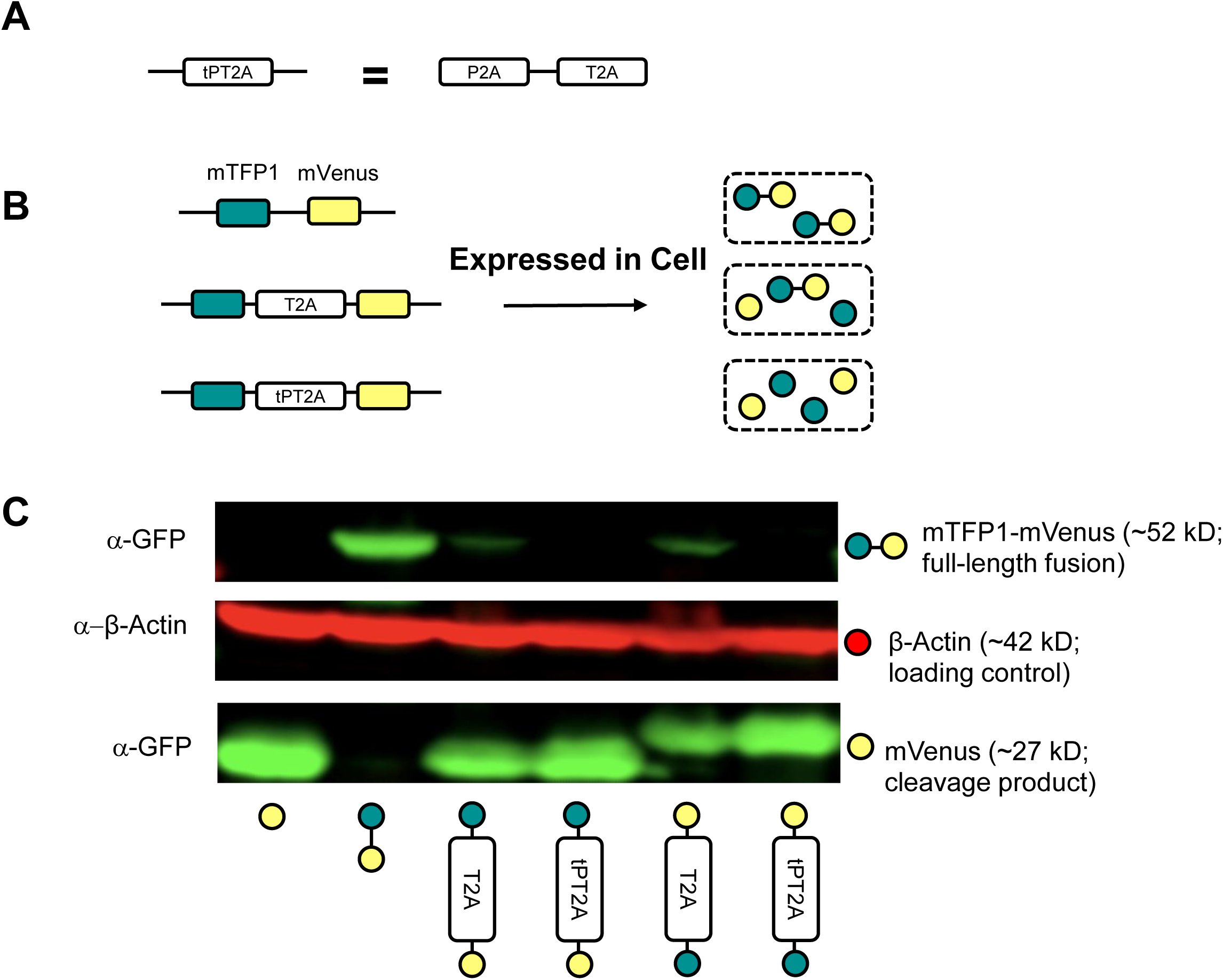
Performance of the Tandem P2A-T2A for Cleaving Barcode Elements. **(A)** tPT2A is a tandem sequence of the P2A and T2A sequences. **(B)** T2A and tPT2A sequences can be used to prevent the fusion of fluorescent proteins expressed by the same plasmid within a cell. **(C)** Western blot comparing the effectiveness of T2A and tPT2A in preventing the fusion of fluorescent proteins in a barcode. HEK293T cells were transfected with indicated plasmids containing variations of mTFP1 and mVenus, then lysates harvested and subjected to western blotting. The anti-GFP antibody recognizes mVenus. Residual fusion protein is not detectable with tPT2A.

**Figure S2.**
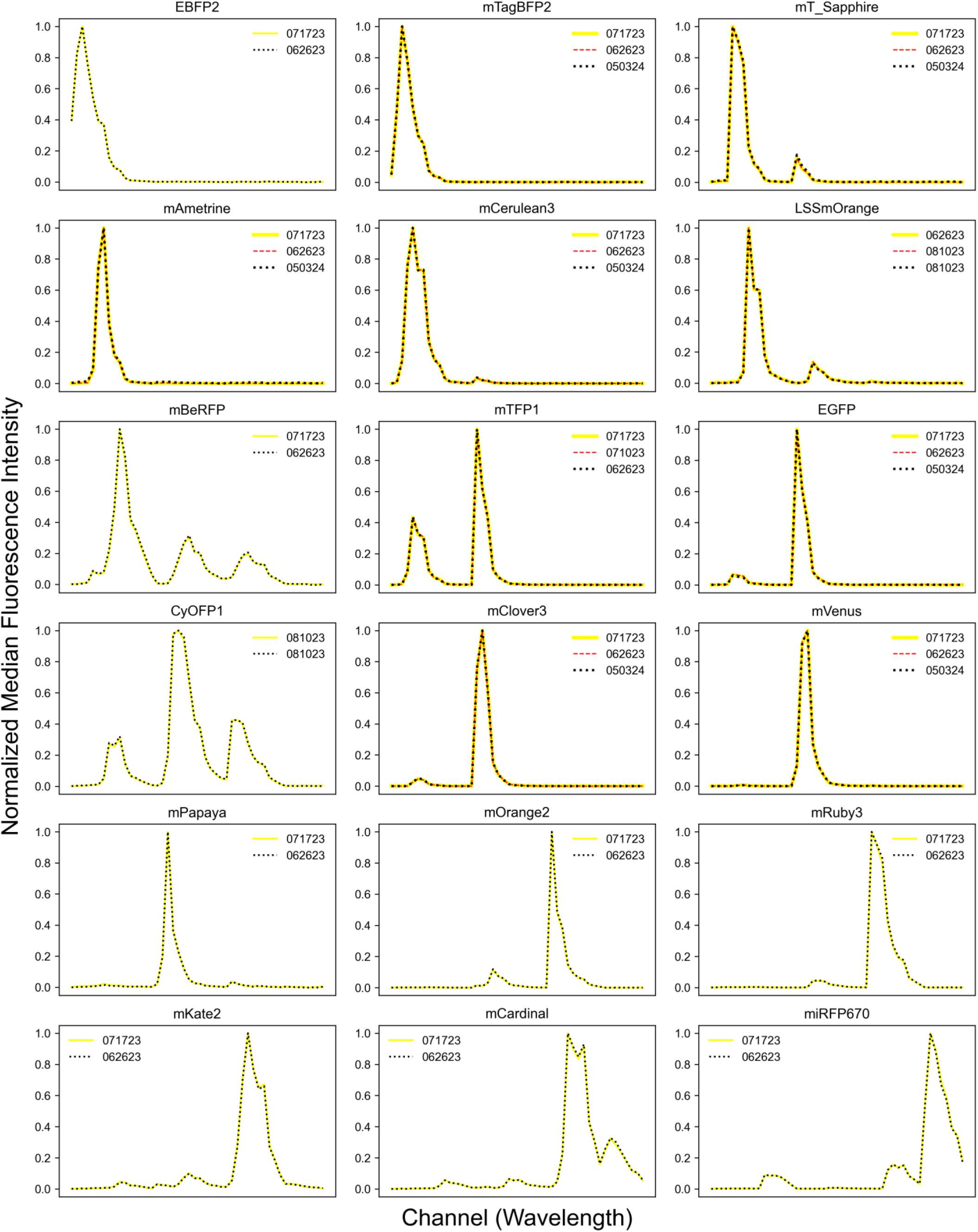
Fluorescent Protein Emission Spectra. HEK293T cells were transfected with plasmids encoding each of the 18 fluorescent proteins(FP), and the normalized median emission intensity spectra were measured using spectral flow cytometry. At least two replicates were performed for each pR-FP probe. In each subplot, different replicates are represented by different colors, with their respective dates used to identify them in the legend.

**Figure S3.**
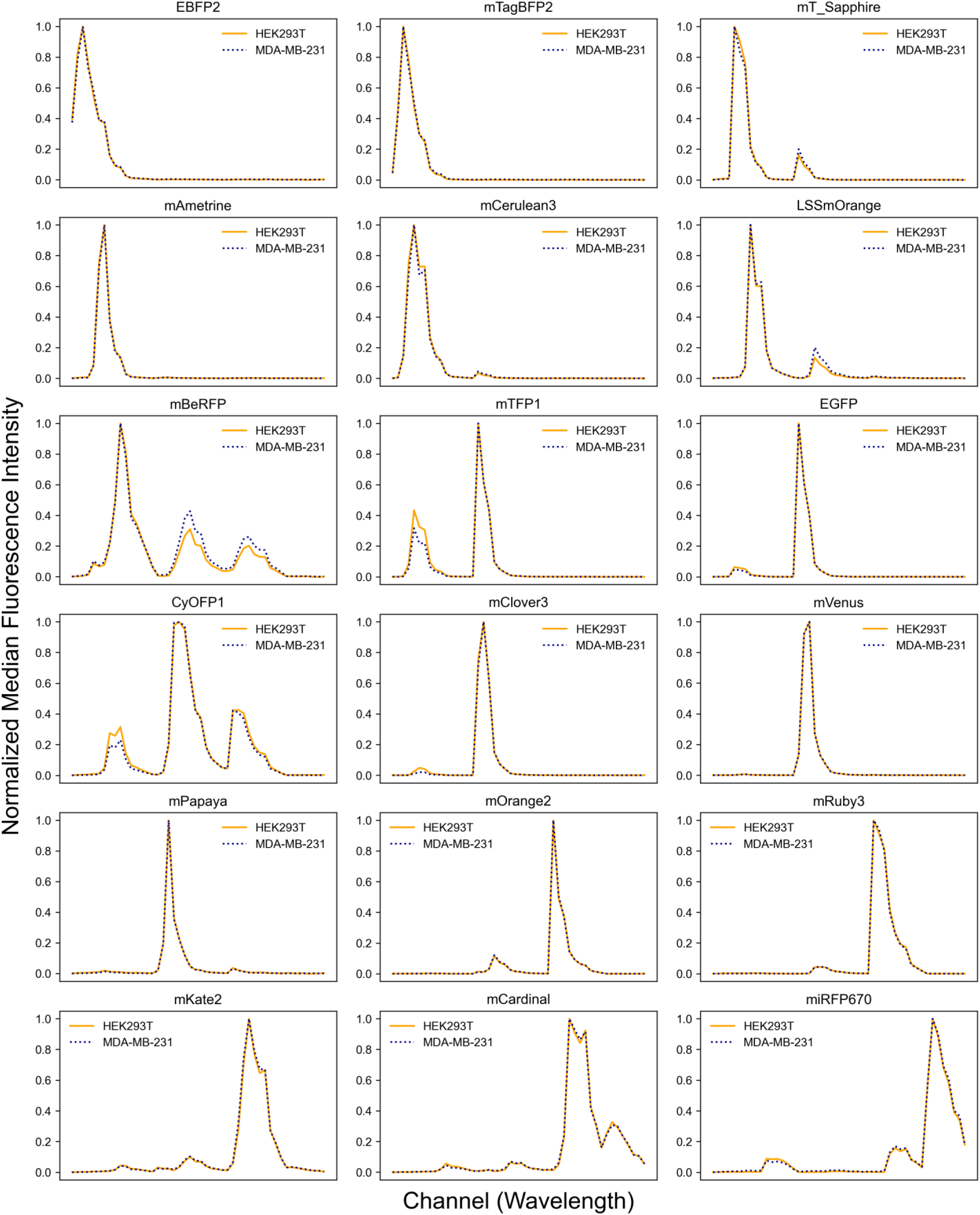
Fluorescent Protein Emission Spectra in HEK293T and MDA-MB-231 Cells. Both cell lines were transfected with plasmids encoding each of the 18 fluorescent proteins, and their normalized median fluorescence intensity spectra were measured using spectral flow cytometry. Each spectra is the average from at least two replicates.

**Figure S4.**
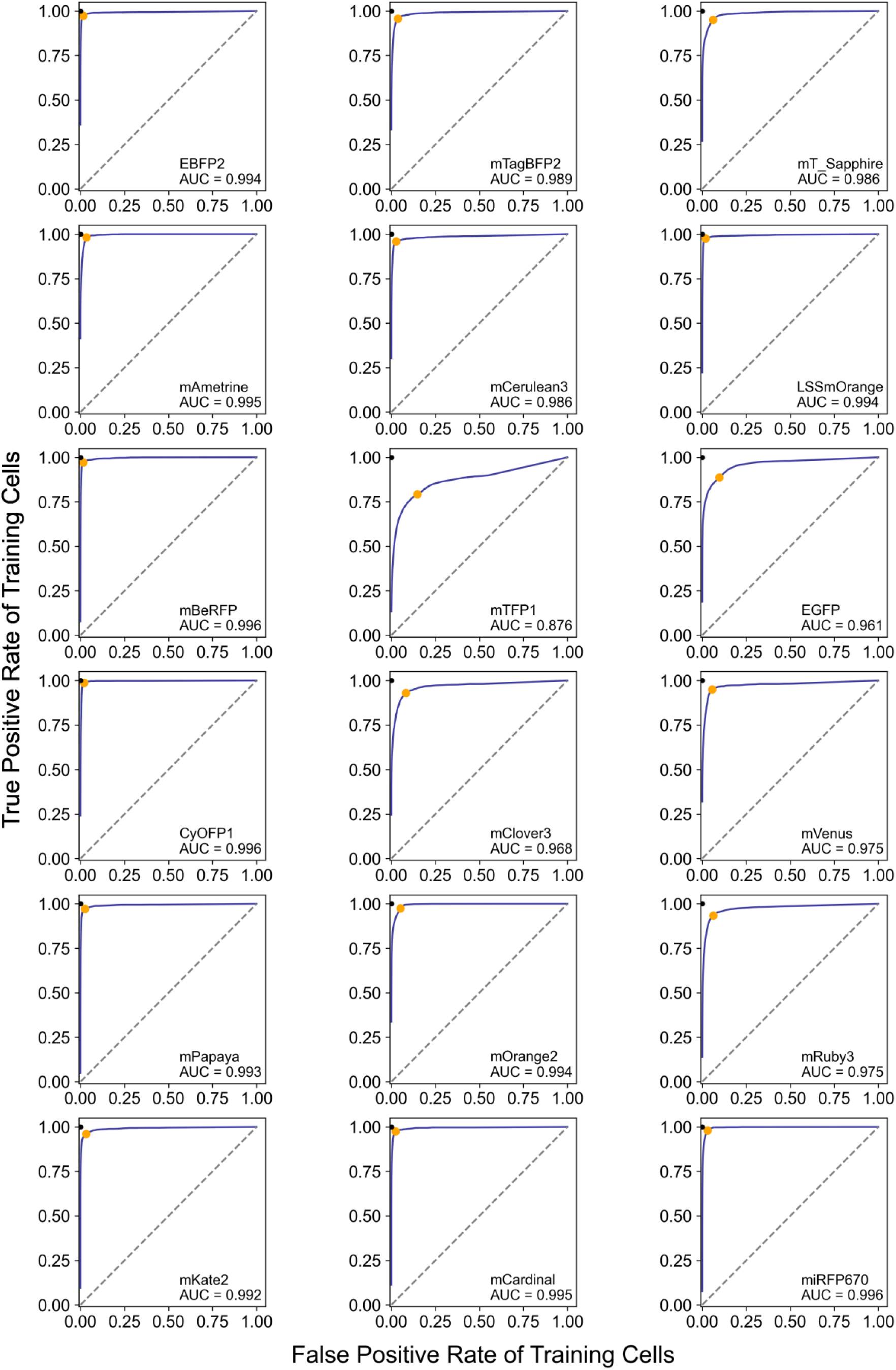
Threshold Estimation for Each Fluorescence Protein from Training Data. For varying thresholds, the training data underwent unmixing, thresholding, and classification to compute the true positive and false positive rates, enabling the construction of a receiver operating characteristic (ROC) curve for each fluorescent protein. The area under the curve (AUC) is displayed in each inset, along with the chosen threshold point (orange) which balances both false positive rate and true positive rate.

**Figure S5.**
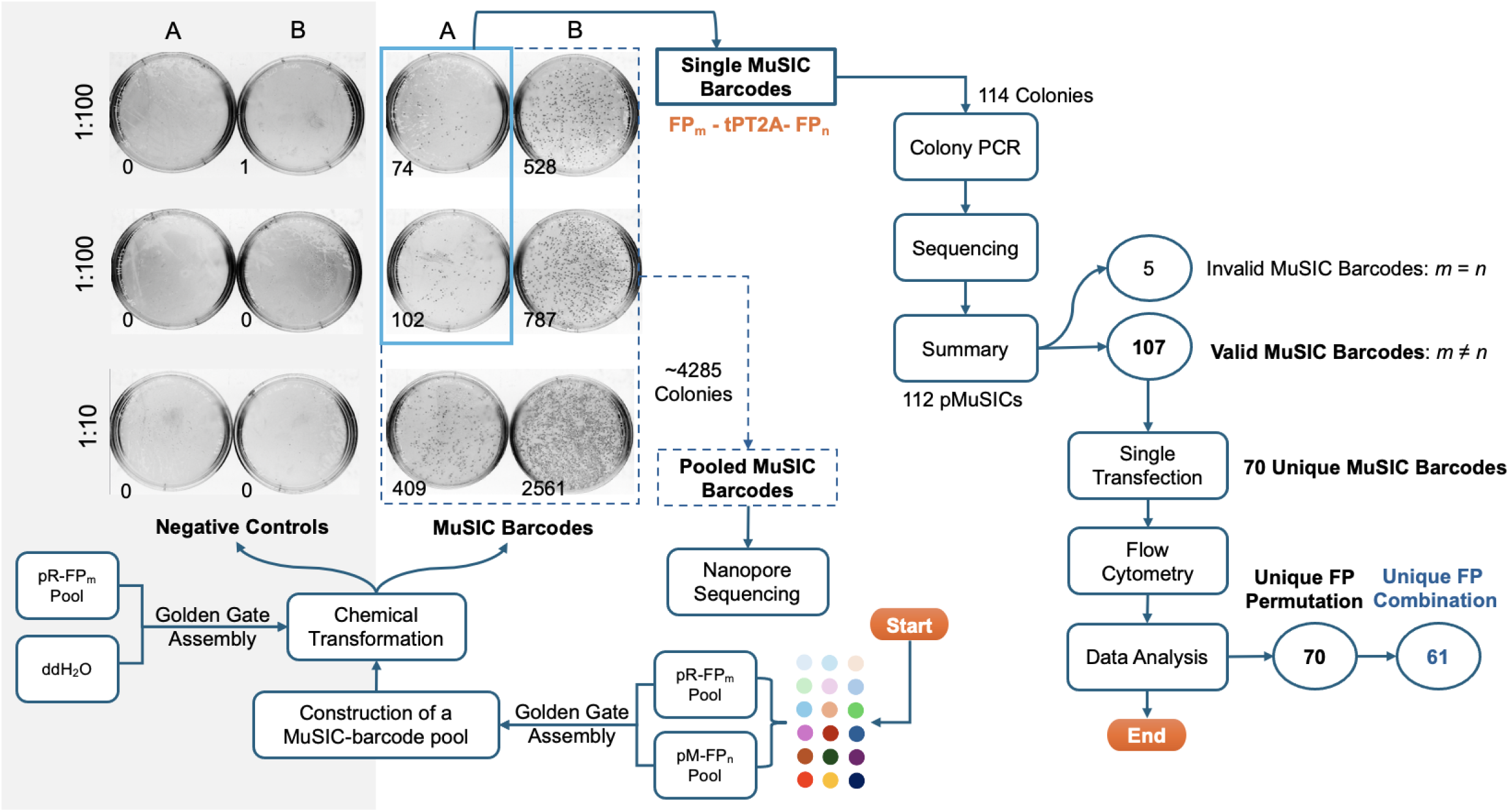
Workflow for Constructing and Analyzing Barcodes. This flowchart outlines the process of constructing the barcode pool through chemical transformation, generating single barcodes for flow cytometry analysis and pooled MuSIC barcodes for nanopore sequencing. To better isolate single colonies, we performed duplicate transformations using a 1:100 dilution of the GoldenGate assembly product. To maximize the colonies without compromising the transformation efficiency, a 1:10 dilution was also performed. We used ddH_2_O in place of the insert pool as the negative control. The number in the bottom left corner of each plate represents the colony count. More than 4000 colonies from the MuSIC-barcode plates, highlighted by the navy dotted square, were scraped and pooled to create the MuSIC barcode library for nanopore sequencing. In the sky blue solid square, 114 colonies were selected and screened by colony PCR and then sequencing (*PlasmidSaurus*). Among 112 positive pMuSICs, 107 contained different FPs and thus were considered valid. Of these, 70 were unique barcodes based on permutation, and 61 were unique combinations.

**Figure S6.**
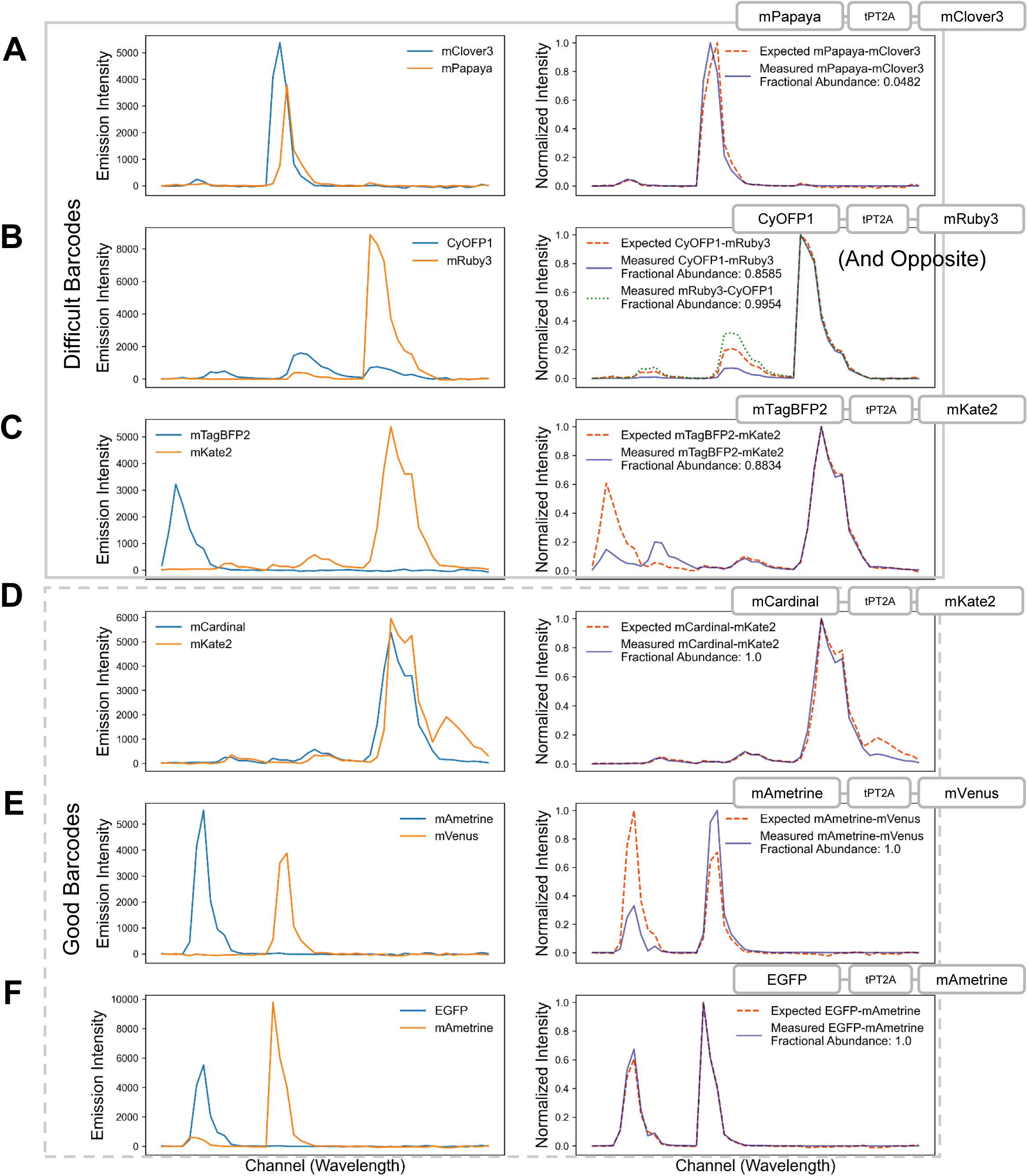
Spectra of Difficult Barcodes (A-C) and Good Barcodes (D-F). (**Left**) Raw intensity median spectra of individual fluorescent proteins (FPs). The raw intensity provides insight into brightness differences (if applicable). (**Right**) Normalized barcode “expected” spectra based on equal expression of the FPs on the left, and experimentally measured spectra. Fractional abundance is the proportion of correctly classified cells. Normalized intensity helps to compare between expected and experimentally measured—brightness differences are built-in. The order of each barcode in the plasmid is depicted at the top right. (**A**) mPapaya-mClover3. (**B**) CyOFP1-mRuby3, and the alternative permutation mRuby3-CyOFP1. (**C**) mTagBFP2-mKate2. (**D**) mCardinal-mKate2. (**E**) mAmetrine-mVenus. (**F**) EGFP-mAmetrine.

**Figure S7.**
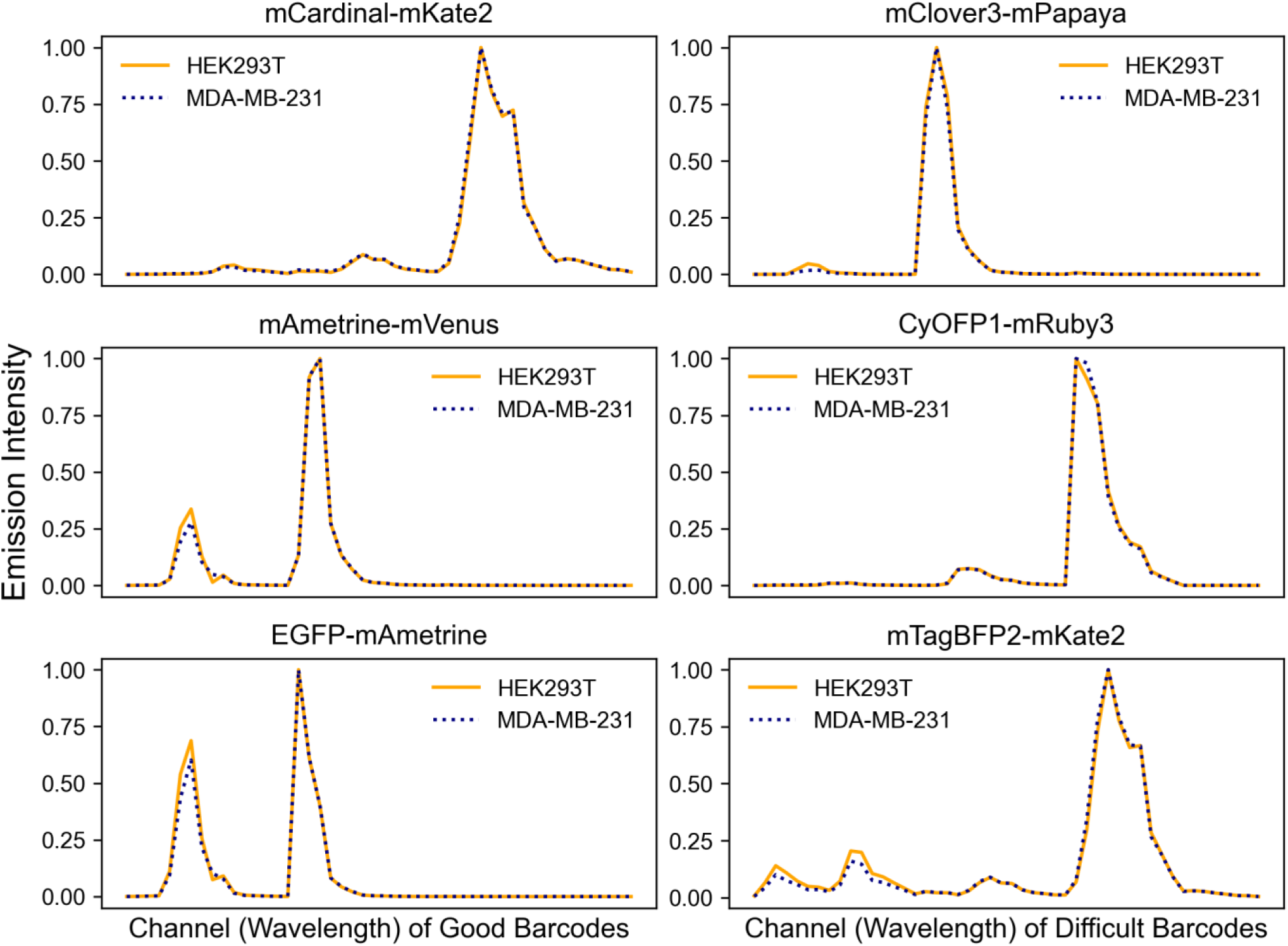
Spectra of Good (Left) or Difficult (Right) Barcodes in Different Cell Lines. Spectra were generated and analyzed as in Figure S6. Briefly, HEK293T or MDA-MB-231 cells were transfected with the barcodes as indicated then analyzed by spectral flow cytometry. Spectra are nearly identical in both cell lines.

**Figure S8.**
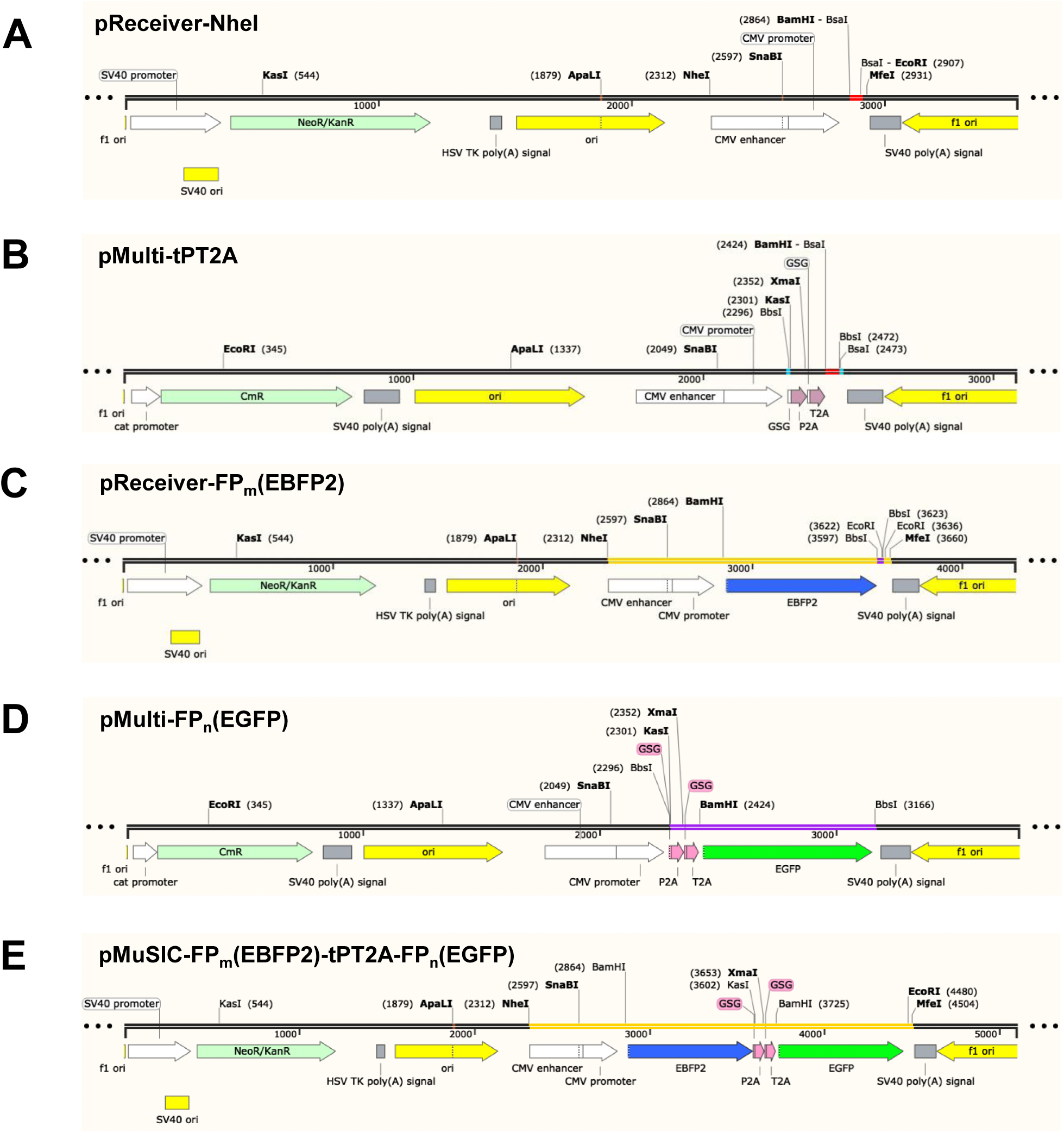
Schematic Representations of Plasmid Backbones, FPs, and a MuSIC Barcode. The schematic plasmid maps were generated using SnapGene. (**A**) An NheI site was inserted upstream of the CMV promoter in pReceiver to create pReceiver-NheI. A pair of BsaI sites (red) was used for insertion of an individual fluorescent protein (FP) as the first probe. (**B**) A pair of BbsI sites (light blue) was introduced upstream of T2A (in pMulti) or P2A-T2A (in pMulti-tPT2A, shown here) and downstream of the BsaI-spacer-BsaI FP insertion cassette (red). This design allowed the insertion of a second FP into pReceiver-FP to generate MuSIC barcodes, while the BsaI sites facilitated individual FP insertion. (**C**) An individual example FP_m_ (EBFP2) was inserted into pReceiver-NheI via BsaI sites (as shown in **A**) to generate pReceiver-FP_m_. A BbsI-TAA-BbsI cassette (purple) was included to terminate transcription for individual testing, and enable the loading of a second FP for MuSIC barcode generation. (**D**) An individual example FP_n_ (EGFP) was inserted into pMulti-tPT2A via BsaI sites (as shown in **B**), generating pMulti-FP_n_. The second FP, containing tPT2A and FP_n_, is highlighted in purple. (**E**) The pMuSIC construct contains FP_m_-tPT2A-FP_n_, exemplified here as EBFP2-tPT2A-EGFP. NheI and MfeI sites in **C** and **E** were used to generate fragments (orange) for nanopore sequencing.

**Figure S9.**
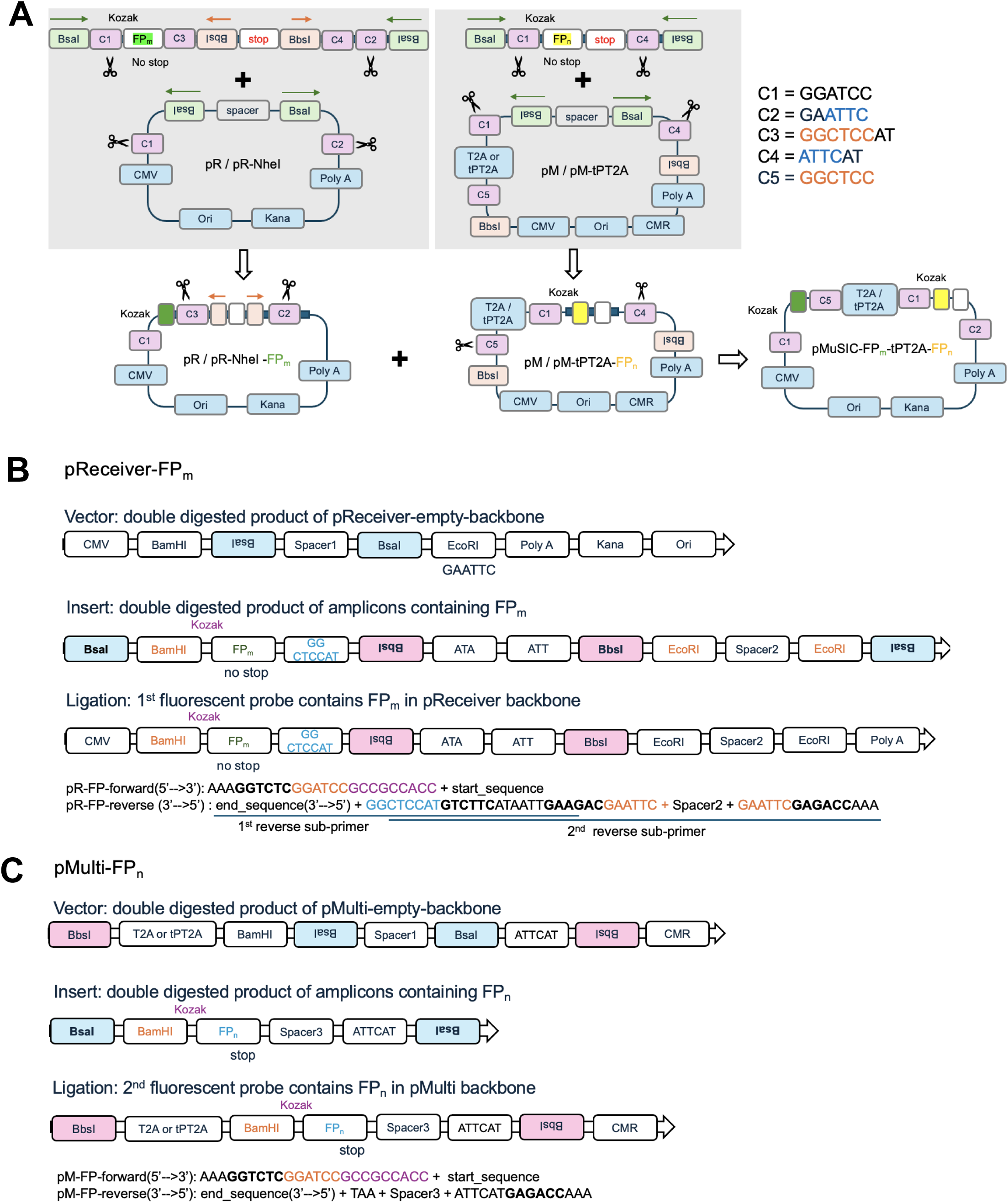
pMuSIC Construction. (**A**) Constructing pMuSIC by Goldengate. First, the BsaI-Spacer-BsaI cassettes in pR were removed by BbsI digestion, and the PCR products containing a fluorescent protein (FP, index *m*) and BbsI-TAA-BbsI cassette (stop), were inserted to create the vector plasmid pR-NheI-FP_m_-BbsI-TAA-BbsI. Similarly, pM underwent BsaI digestion for the FP insertion (FP, index *n*), resulting in the insert plasmids pM-FP_n_. Finally, both vector and insert plasmids were digested at BbsI sites to generate pMuSICs containing the MuSIC barcode (FP_m_-2A-FP_n_). (**B-C**) Primer designs to amplify the FP inserts for both the vector and insert plasmids, respectively. The sequences of all 18 FPs were used as templates (**Table S5**), with their start and end sequences listed in **Table S6**, along with the sequences of the spacers.

**Figure S10.**
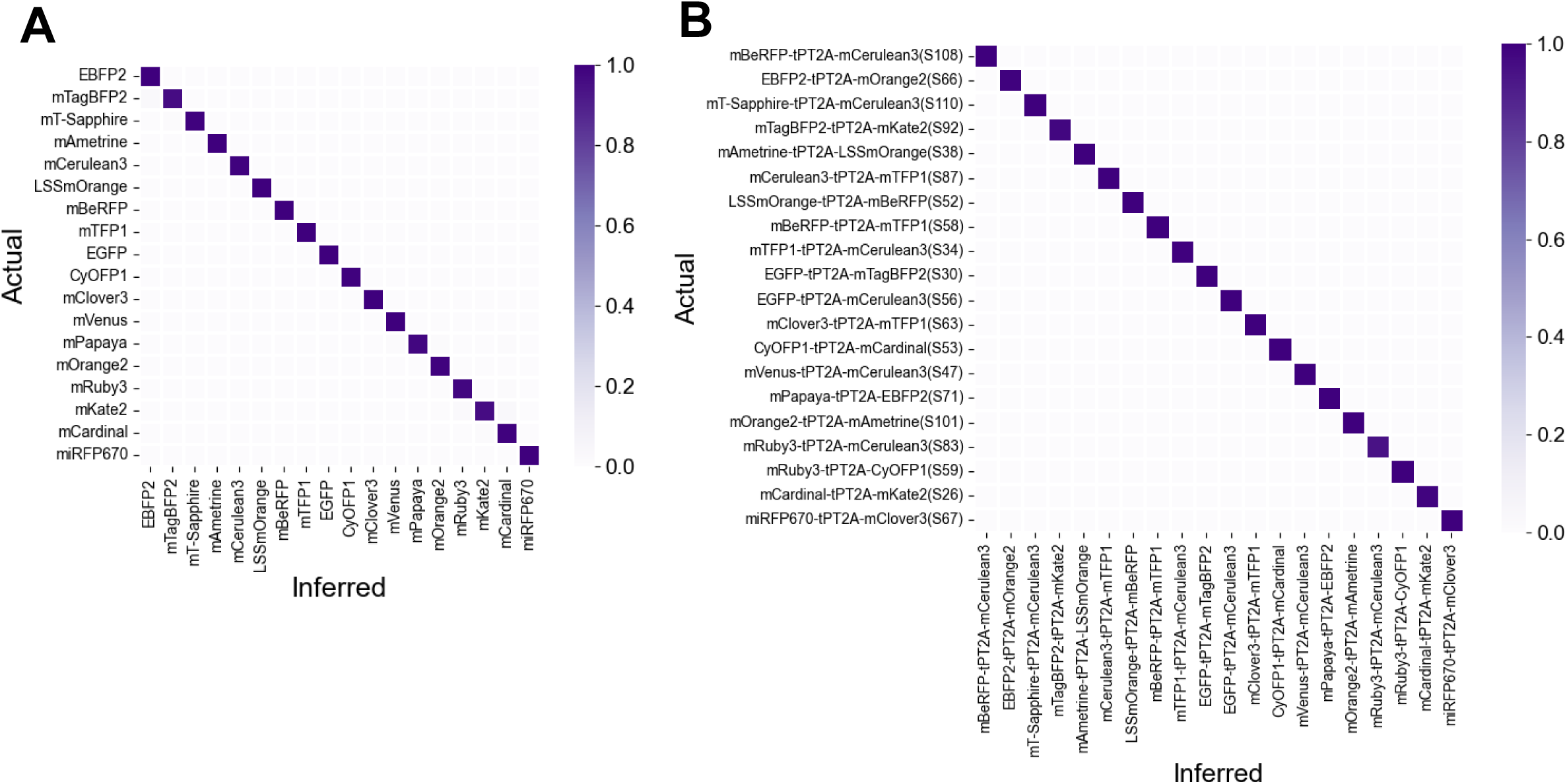
Nanopore Sequencing Validation Results. Classification of (**A**) 18 pR-FPs and (**B**) 20 barcodes Color bar denotes fraction inferred.

